# Evolutionary conservation of centriole rotational asymmetry in the human centrosome

**DOI:** 10.1101/2021.07.21.453218

**Authors:** Noémie Gaudin, Paula Martin Gil, Meriem Boumendjel, Dmitry Ershov, Catherine Pioche-Durieu, Manon Bouix, Quentin Delobelle, Lucia Maniscalco, Thanh Bich Ngan Phan, Vincent Heyer, Bernardo Reina-San-Martin, Juliette Azimzadeh

**Affiliations:** Université de Paris, CNRS, Institut Jacques Monod, 75013, Paris, France.; Image Analysis Hub, C2RT, Institut Pasteur, Paris, France.; Hub de Bioinformatique et Biostatistique – Département Biologie Computationnelle, Institut Pasteur, USR 3756 CNRS, Paris, France.; Institut de Génétique et de Biologie Moléculaire et Cellulaire (IGBMC), Illkirch, France.; Institut National de la Santé et de la Recherche Médicale (INSERM), U1258, Illkirch, France.; Centre National de la Recherche Scientifique (CNRS), UMR7104, Illkirch, France.; Université de Strasbourg, Illkirch, France.

**Keywords:** centriole, centrosome, LRRCC1, VFL1, C2CD3, Joubert syndrome, asymmetry.

## Abstract

Centrioles are formed by microtubule triplets in a nine-fold symmetric arrangement. In flagellated protists and in animal multiciliated cells, accessory structures tethered to specific triplets render the centrioles rotationally asymmetric, a property that is key to cytoskeletal and cellular organization in these contexts. In contrast, centrioles within the centrosome of animal cells display no conspicuous rotational asymmetry. Here, we uncover rotationally asymmetric molecular features in human centrioles. Using ultrastructure expansion microscopy, we show that LRRCC1, the ortholog of a protein originally characterized in flagellate green algae, associates preferentially to two consecutive triplets in the distal lumen of human centrioles. LRRCC1 partially co-localizes and affects the recruitment of another distal component, C2CD3, which also has an asymmetric localization pattern in the centriole lumen. Together, LRRCC1 and C2CD3 delineate a structure reminiscent of a filamentous density observed by electron microscopy in flagellates, termed the ‘acorn’. Functionally, the depletion of LRRCC1 in human cells induced defects in centriole structure, ciliary assembly and ciliary signaling, supporting that LRRCC1 cooperates with C2CD3 to organizing the distal region of centrioles. Since a mutation in the *LRRCC1* gene has been identified in Joubert syndrome patients, this finding is relevant in the context of human ciliopathies. Taken together, our results demonstrate that rotational asymmetry is an ancient property of centrioles that is broadly conserved in human cells. Our work also reveals that asymmetrically localized proteins are key for primary ciliogenesis and ciliary signaling in human cells.

## Introduction

Centrioles are cylindrical structures with a characteristic ninefold symmetry, which results from the arrangement of their constituent microtubule triplets (LeGuennec et al., 2021). In animal cells, centrioles are essential for the assembly of centrosomes and cilia. The centrosome, composed of two centrioles embedded in a pericentriolar material (PCM), is a major organizer of the microtubule cytoskeleton. In addition, most vertebrate cells possess a primary cilium, a sensory organelle that assembles from the oldest centriole within the centrosome, called mother centriole (Kumar and Reiter, 2021).

Centrioles within the centrosome show no apparent rotational asymmetry, *i.e.,* no structural asymmetry of the microtubule triplets. In vertebrates, the mother centriole carries distal appendages (DAs) and subdistal appendages arranged in a symmetric manner around the centriole cylinder (Kumar and Reiter, 2021). In contrast, the centriole/basal body complex of flagellates, to which the animal centrosome is evolutionary related, is characterized by marked rotational asymmetries (Azimzadeh, 2021; Yubuki and Leander, 2013). In flagellates, an array of fibers and microtubules anchored asymmetrically at centrioles controls the spatial organization of the cell (Feldman et al., 2007; Yubuki and Leander, 2013). The asymmetric attachment of cytoskeletal elements appears to rely on molecular differences between microtubule triplets. In the green alga *Chlamydomonas reinhardtii,* Vfl1p (Variable Flagella number 1 protein) localizes principally at two triplets near the attachment site of a striated fiber connecting the centrioles (Silflow et al., 2001). This fiber is absent or mispositioned in the *vfl1* mutant, leading to defects in centriole position and number, and overall cytoskeleton disorganization (Adams et al., 1985; Feldman et al., 2007). In the same region, a rotationally asymmetric structure termed the ‘acorn’ was observed in the centriole lumen by transmission electron microscopy. The acorn appears as a filament connecting five successive triplets and is in part colocalized with Vfl1p (Geimer and Melkonian, 2005, 2004).

We recently established that Vfl1p function is conserved in the multiciliated cells (MCCs) of planarian flatworms, which was recently confirmed in xenopus (Basquin et al., 2019; Nommick et al., 2022). MCCs assemble large numbers of centrioles that are polarized in the plane of the plasma membrane to enable the directional beating of cilia (Meunier and Azimzadeh, 2016), like in *C. reinhardtii*. The planarian ortholog of Vfl1p is required for the assembly of two appendages that decorate MCC centrioles asymmetrically, the basal foot and the ciliary rootlet (Basquin et al., 2019). Depleting Vfl1p orthologs in planarian or xenopus MCCs alters centriole rotational polarity, reminiscent of the *vfl1* phenotype in *C. reinhardtii* (Adams et al., 1985; Basquin et al., 2019; Nommick et al., 2022). Intriguingly, the human ortholog of Vfl1p, called LRRCC1 (Leucine Rich Repeat and Coiled Coil containing 1) localizes at the centrosome despite the lack of rotationally asymmetric appendage in this organelle (Andersen et al., 2003; Muto et al., 2008). Furthermore, a homozygous mutation in the *LRRCC1* gene was identified in two siblings affected by a ciliopathy called Joubert syndrome (JBTS), suggesting that LRRCC1 might somehow affect the function of non-motile cilia (Shaheen et al., 2016).

Here, we show that LRRCC1 localizes in a rotationally asymmetric manner in the centrioles of the human centrosome. We further establish that LRRCC1 is required for proper ciliary assembly and signaling, which likely explains its implication in JBTS. LRRCC1 affects the recruitment at centrioles of another ciliopathy protein called C2CD3 (C2 domain containing 3), which we found to also localize in a rotationally asymmetric manner, forming a pattern partly reminiscent of the acorn described in flagellates. Our findings uncover the unanticipated rotational asymmetry of centrioles in the human centrosome and show that this property is connected to the assembly and function of primary cilia.

## Results

### LRRCC1 localizes asymmetrically at the distal end of centrioles

To investigate a potential role of LRRCC1 at the centrosome, we first sought to determine its precise localization. We raised antibodies against two different fragments within the long C-terminal coiled-coil domain of LRRCC1 (Ab1, 2), which both stained the centrosome region in human Retinal Pigmented Epithelial (RPE1) cells (Fig. 1a; Supplemental Fig. S1a), as previously reported (Muto et al., 2008). Labeling intensity was decreased in LRRCC1-depleted cells for both antibodies, supporting their specificity (Fig. 4a, b; Supplemental Fig. S1d, e, g). LRRCC1 punctate labeling in the centrosomal region indicated that it is present within centriolar satellites, confirming a previous finding that LRRCC1 interacts with the satellite component PCM1 (Gupta et al., 2015). After nocodazole depolymerization of microtubules to disperse satellites, a fraction of LRRCC1 was retained at centrioles (Fig. 1a; Supplemental Fig. S1a), providing evidence that LRRCC1 is also a core component of centrioles. To determine LRRCC1 localization more precisely within the centriolar structure, we used ultrastructure expansion microscopy (U-ExM) (Gambarotto et al., 2019) combined with imaging on a Zeiss Airyscan 2 confocal microscope, thereby increasing the resolution by a factor of ∼ 8 compared to conventional confocal microscopy. We found that LRRCC1 localizes at the distal end of centrioles as well as of procentrioles (Fig. 1b). Strikingly, and unlike other known centrosome components, LRRCC1 decorated the distal end of centrioles in a rotationally asymmetric manner. Indeed, LRRCC1 was detected close to the triplet blades and towards the lumen of the centriole (Fig. 1c). The staining was often associated with two or more consecutive triplets, one of them being usually more brightly labelled than the others. In addition, a fainter staining was consistently detected along the entire length of all triplets (Fig. 1b, brighter exposure). This pattern was observed in both RPE1 and HEK 293 cells and was obtained with both anti-LRRCC1 antibodies (Supplemental Fig. S1h), supporting its specificity. We verified that LRRCC1 asymmetric localization was also observed in unexpanded cells by directly analyzing immunofluorescence samples by Airyscan microscopy (Fig. 1d). We measured the lateral distribution of LRRCC1 signal intensity peak relative to the long axis of the centriole. The distance between peaks was greater for LRRCC1 than for hPOC5, a marker that localizes symmetrically in the centriole (Azimzadeh et al., 2009; le Guennec et al., 2020), confirming the asymmetry of LRRCC1 staining. The distal pattern obtained by U-ExM showed some variability, especially in the distance between LRRCC1 and the centriole wall (Fig. 1c), which could result from the fact that centrioles were not perfectly orthogonal to the imaging plan. To obtain a more accurate picture of LRRCC1 localization, we generated 3D reconstructions that we realigned, first along the vertical axis, then with respect to one another using the most intense region of the LRRCC1 labeling as a reference point (Fig. 1e; Supplemental Fig. S2a- b). An average 3D reconstruction was then generated (Fig. 1f) and revealed that LRRCC1 was mainly associated to one triplet, and to a lesser extent to its direct neighbor counterclockwise, on their luminal side. A longitudinal view confirmed that LRRCC1 is principally located at the distal end of centrioles.

**Figure 1.**
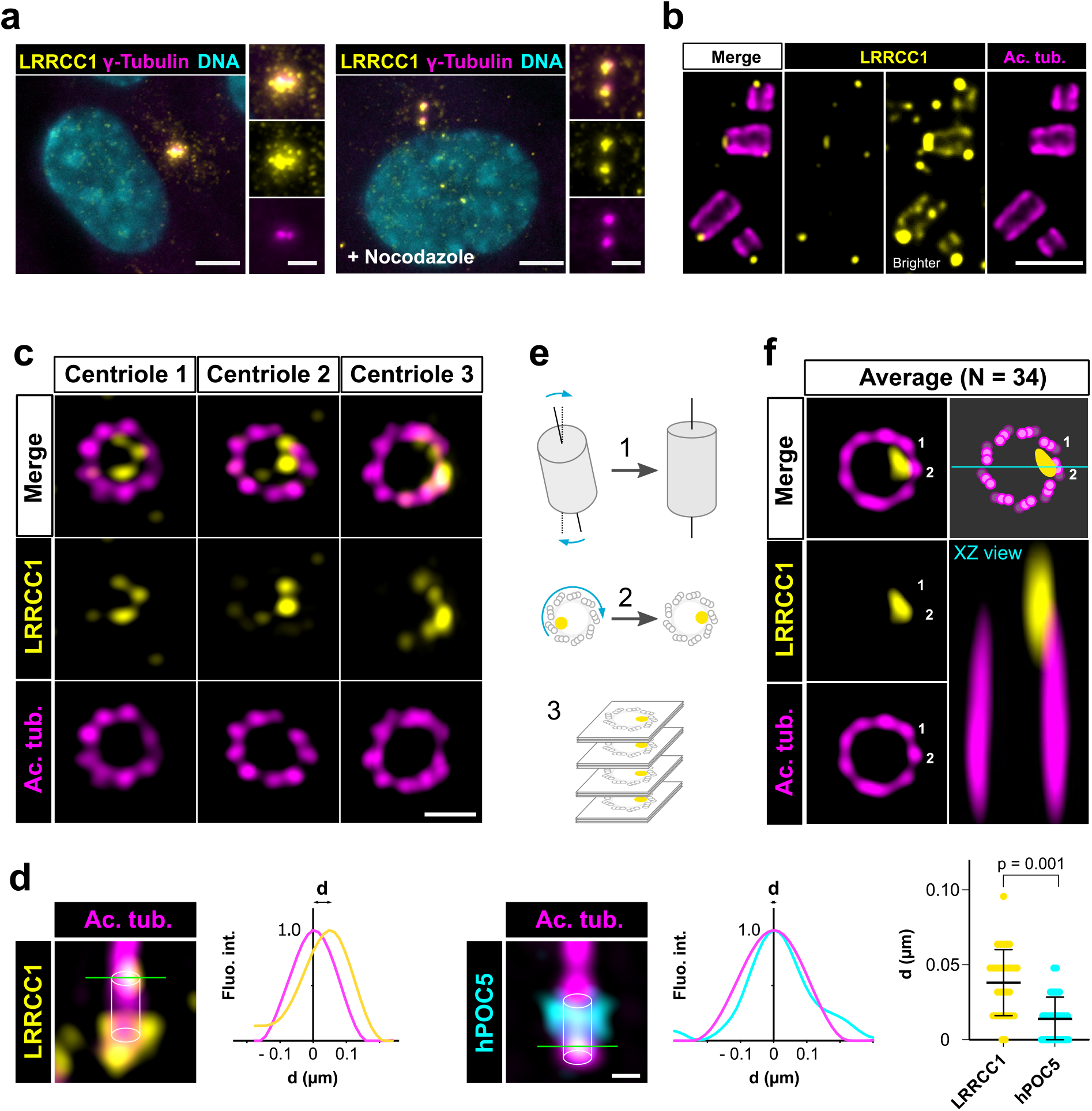
LRRCC1 is localized in a rotationally asymmetric manner at the distal end of centrioles in the human centrosome. **a)** LRRCC1 localization in non-treated RPE1 cells (left) or in cells treated with nocodazole to disperse the pericentriolar satellites (right). LRRCC1 (Ab2, yellow), γ-tubulin (PCM, magenta) and DNA (cyan). Bar, 5 µm (insets, 2 µm). **b)** Longitudinal view of centrioles and procentrioles in the duplicating centrosome of an RPE1 cell analyzed by U-ExM. LRRCC1 (Ab2, yellow), acetylated tubulin (magenta). Bar, 0.5 µm. **c)** Centrioles from WT RPE1 cells as seen from the distal end. LRRCC1 (Ab2, yellow), acetylated tubulin (magenta). Images are maximum intensity projections of individual z-sections encompassing the LRRCC1 signal. Note that an apparent shift between channels occurs when centrioles are slightly angled with respect to the imaging axis. Bar, 0.2 µm. **d)** Lateral distance between LRRCC1 (left, yellow) or hPOC5 (middle, cyan) signal intensity peaks and the centriole center (given by the position of acetylated tubulin intensity peak, magenta) in ciliated RPE1 cells. Individual intensity profiles were measured along the green lines. The approximate position of the centriole is shown (white cylinders). Note that LRRCC1 and hPOC5 were also detected at the periphery of the centriole. Right: interpeak distance (d). Bars, mean ± SD, 31 cells from 2 different experiments (Kolmogorov-Smirnov test). **e)** Workflow for calculating the average staining from 3D-reconstructed individual centrioles generated from confocal z-stacks. The brightest part of LRRCC1 signal was used as a reference point to align the centrioles. **f)** Average LRRCC1 staining obtained from 34 individual centrioles viewed from the distal end, in transverse and longitudinal views. A diagram representing the average pattern in transverse view is also shown.

Together, our results show that LRRCC1 is localized asymmetrically within the distal centriole lumen, establishing that centrioles within the human centrosome are rotationally asymmetric.

### The localization pattern of LRRCC1 is similar at the centrosome and in mouse MCCs

LRRCC1 orthologs are required for establishing centriole rotational polarity in planarian and xenopus MCCs, like in *C. reinhardtii* (Basquin et al., 2019; Nommick et al., 2022; Silflow et al., 2001). It is therefore plausible that LRRCC1-related proteins localize asymmetrically in MCC centrioles, and indeed, Lrrcc1 was recently found associated to the ciliary rootlet in xenopus MCCs (Nommick et al., 2022). To determine whether LRRCC1 also localizes at the distal end of MCC centrioles in addition to its rootlet localization, and if so, whether LRRCC1 localization pattern resembles that observed at the centrosome, we analyzed mouse ependymal and tracheal cells by U-ExM. In *in vitro* differentiated ependymal cells, the labeling generated by the anti-LRRCC1 antibody was consistent with our observations in human culture cells. Mouse Lrrcc1 localized asymmetrically at the distal end of centrioles, opposite to the side where the basal foot is attached (Fig. 2a), as determined by co-staining with the basal foot marker γ-tubulin (Clare et al., 2014). Lrrcc1 was also present at the distal end of procentrioles forming via either the centriolar or acentriolar pathways (*i.e.,* around parent centrioles or deuterosomes, respectively) (Fig. 2b). We also examined tracheal explants, in which centrioles were docked and polarized at the apical membrane in higher proportions (Fig. 2c). We obtained an average image of Lrrcc1 labeling from 35 individual centrioles aligned using the position of the basal foot as a reference point. This revealed that Lrrcc1 is principally located in the vicinity of 3 triplets opposite to the basal foot, to the right of basal foot main axis (triplet number 9, 1 and 2 on the diagram in Fig. 2d). Lrrcc1 was located farther away from the triplet wall than in centrioles of the centrosome, but this was likely an effect of a deformation of the centrioles (Fig. 2c, d) caused by the incomplete expansion of the underlying cartilage layer in tracheal explants. In agreement, Lrrcc1 was close to the triplets in ependymal cell monolayers, which expand isometrically. Besides the distal centriole staining, we found no evidence that Lrrcc1 is associated to the ciliary rootlet in mouse MCCs, unlike in xenopus. The Lrrcc1 pattern in mouse MCCs was thus similar to the pattern observed at the human centrosome.

**Figure 2.**
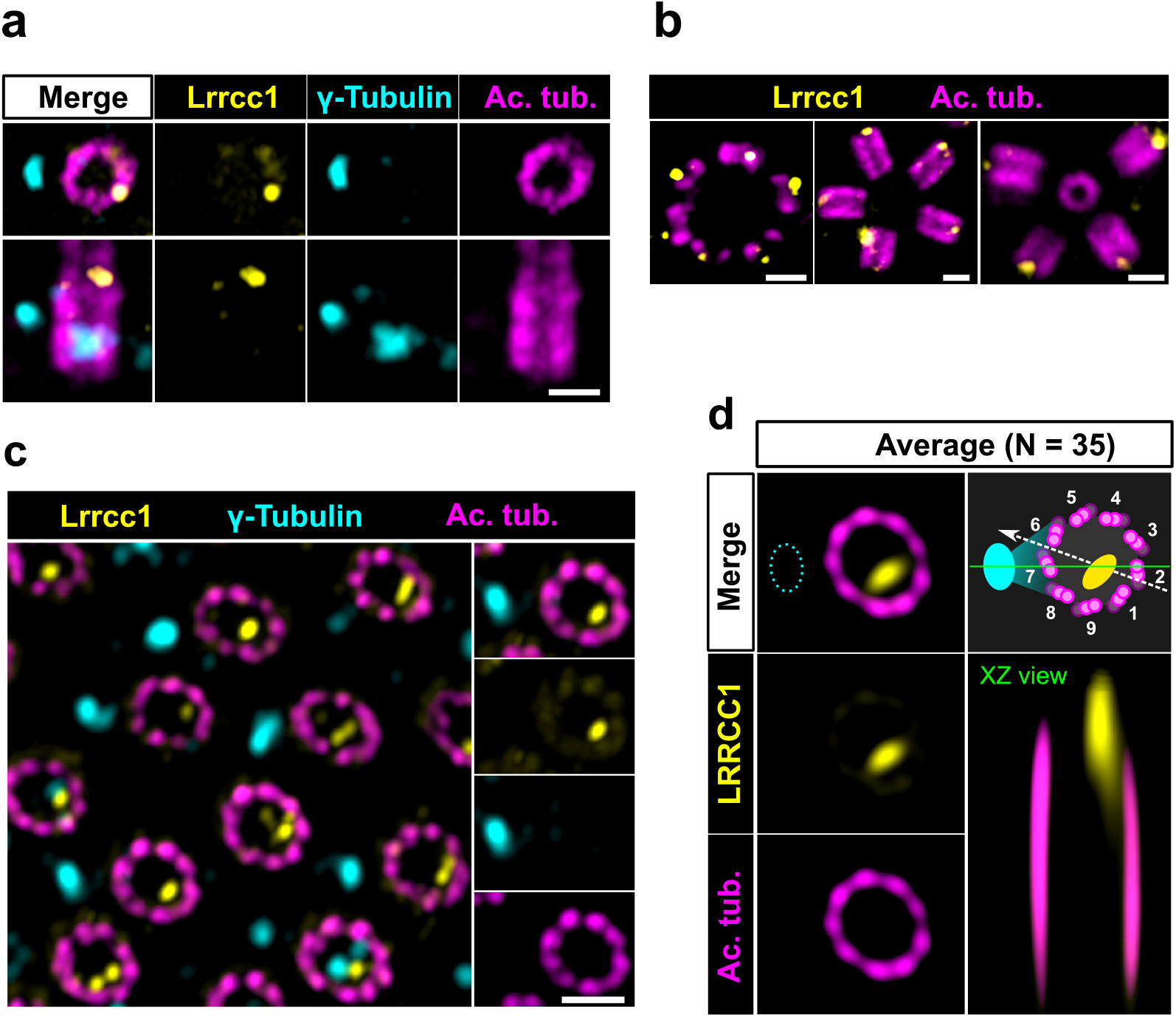
The LRRCC1 rotationally asymmetric pattern is conserved in mouse MCCs. **a)** Centrioles in the cytoplasm of mouse ependymal cells differentiating *in vitro* analyzed by U-ExM, in longitudinal and transverse view. Lrrcc1 (Ab2, yellow), γ-tubulin (basal foot cap, cyan) and acetylated tubulin (magenta). Of note, γ-tubulin was also detected in the proximal lumen of centrioles. Bar, 0.2 µm. **b)** Procentrioles assembling via the centriolar (right) or the deuterosome pathway (left and center) in ependymal cells. Lrrcc1 (Ab2, yellow), acetylated tubulin (magenta). Bar, 0.2 µm. **c)** Transverse view of centrioles docked at the apical membrane in fully differentiated mouse tracheal cells, viewed from the distal end. Lrrcc1 (Ab2, yellow), γ-tubulin (cyan) and acetylated tubulin (magenta). Bar, 0.2 µm. **d)** Average image generated from 35 individual centrioles from mouse trachea, viewed from the distal end, shown in transverse and longitudinal views. The position of the basal foot (cyan dotted line) stained with γ-tubulin was used as a reference point to align the centrioles. A diagram of the average pattern in transverse view is shown, in which the direction of ciliary beat (Schneiter et al., 2021) is represented by a dotted arrow and the basal foot axis by a green line. Triplets are numbered counterclockwise from the LRRCC1 signal.

Together, these results show that LRRCC1 asymmetric localization is a conserved feature of mammalian centrioles, presumably linked to the control of centriole rotational polarity and ciliary beat direction in MCCs.

### Procentriole assembly site is partly correlated with centriole rotational polarity

In *C. reinhardtii*, cytoskeleton organization and flagellar beat direction depend on the position and orientation at which new centrioles arise during cell division. Reflecting the stereotypical organization of centrioles and procentrioles in this species, Vfl1p is recruited early and at a fixed position at the distal end of procentrioles (Fig.3a) (Geimer and Melkonian, 2004; Silflow et al., 2001). We therefore wondered whether this mechanism might be to some extent conserved at the centrosome, which could explain the persistence of centriole rotational asymmetry despite the absence of asymmetric appendages or ciliary motility in most animal cell types. We first analyzed the timing of LRRCC1 incorporation into procentrioles. LRRCC1 was already present at an early stage of centriole assembly, when the procentrioles stained with acetylated tubulin and the cartwheel component SAS-6 were only about 100 nm in length (Fig. 3b). LRRCC1 was then detected during successive stages of procentriole elongation, always localizing asymmetrically and distally (Fig. 3c), like in *C. reinhardtii*. We then examined LRRCC1 localization in duplicating centrosomes by generating 3D-reconstructions of diplosomes (*i.e.,* orthogonal centriole pairs) from RPE1 and HEK 293 cells processed by U-ExM (Fig. 3d). We analyzed two parameters: the angle between LRRCC1 in the procentriole and the long axis of the parent centriole used as reference (Fig. 3d, LRRCC1 localization in procentrioles), and the angle between procentriole position and LRRCC1 in the parent centriole (Fig. 3d, Procentriole position with respect to centriolar LRRCC1). We found that LRRCC1 localization in procentrioles was more often aligned with the long axis of the parent centriole in RPE1 cells (Fig. 3d, top left panel, quadrants Q1 and Q3, respectively), but less so in HEK 293 cells (top right panel), in which the distribution was closer to a random distribution. Thus, human procentrioles do not arise in a fixed orientation, although there appears to be a bias toward alignment of LRRCC1 with the main axis of the parent centriole in RPE1 cells. Next, we analyzed the position of procentrioles with respect to centriolar LRRCC1 (bottom panels).

**Figure 3.**
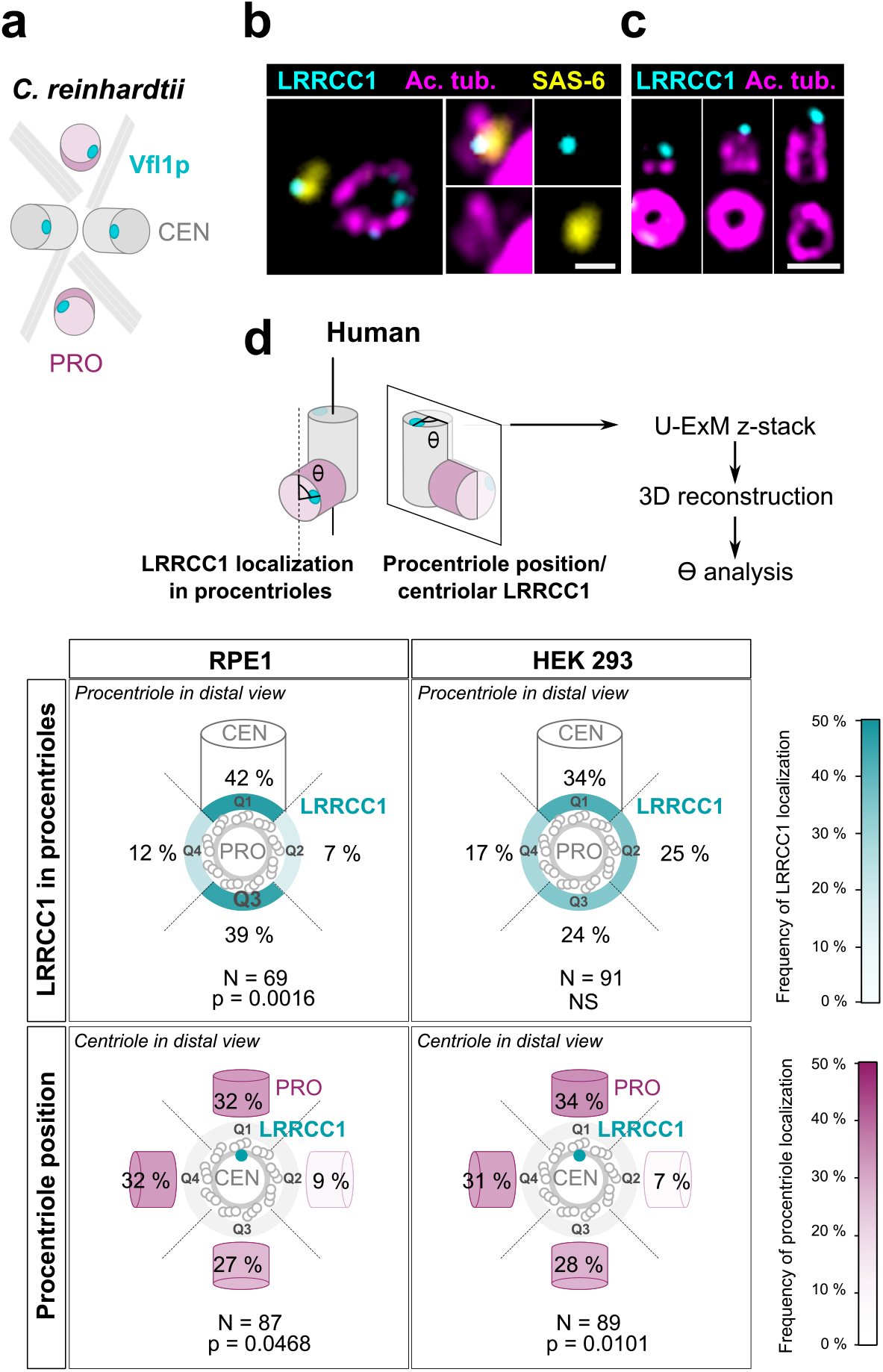
Procentriole assembly site is partly correlated with centriole rotational polarity. **a)** Diagram showing the localization of Vfl1p (cyan) in the centrioles/basal bodies (grey) and procentrioles/probasal bodies (pink) of *C. reinhardtii*. The microtubule roots are also shown. **b)** Early stage of procentriole assembly stained for LRRCC1 (Ab2, cyan), SAS-6 (yellow) and acetylated tubulin (magenta) in a HEK 293 cell. The brightness of the acetylated tubulin labeling was increased in the inserts. Bar, 0.1 µm. **c)** Successive stages of centriole elongation in HEK 293 cells stained for LRRCC1 (Ab2, cyan) and acetylated tubulin (magenta). Bar, 0.1 µm. **d)** Location of LRRCC1 in the procentrioles (top panels) and position of the procentriole relative to its parent centriole polarity (bottom panels), in RPE1 and HEK 293 centrioles analyzed by U-ExM. For each diplosome, the angle between LRRCC1 in the procentriole and the centriole long axis (top panels), or between the procentriole and LRRCC1 in the centriole (bottom panels) was measured. The number of diplosomes analyzed is indicated. p values are indicated when statistically different from a random distribution (χ^2^-test).

Based on current models, procentriole assembly is expected to occur at a random position around parent centrioles in animal cells (Takao et al., 2019). Identification of LRRCC1 provided the first opportunity to directly test this model. In diplosomes from both RPE1 and HEK 293 cells, the position of procentrioles with respect to LRRCC1 location in the parent centriole was variable, confirming that the position at which procentrioles assemble is not strictly controlled in human cells. Interestingly, however, the procentrioles were not distributed in a completely random fashion either. Procentrioles were found in quadrant Q2 (45-135 degrees clockwise from LRRCC1 centroid) on average 4 times less often than in the other quadrants, both in RPE1 and HEK 293 cells, suggesting that rotational polarity of the parent centriole somehow impacts procentriole assembly.

Overall, these results suggest that centriole rotational polarity influences centriole duplication, limiting procentriole assembly within a particular region of centriole periphery. Nevertheless, procentrioles are not formed at a strictly determined position, suggesting that the mechanisms involving the LRRCC1 ortholog Vfl1p in centriole duplication in *C. reinhardtii* are not or not completely conserved at the centrosome.

### LRRCC1 is required for primary cilium assembly and ciliary signaling

A previous report identified a homozygous mutation in a splice acceptor site of the *LRRCC1* gene in two siblings diagnosed with JBTS (Shaheen et al., 2016), but how disruption of LRRCC1 expression affects ciliary assembly and signaling has never been investigated. To address this, we generated RPE1 cell lines deficient in LRRCC1 using two different CRISPR/Cas9 strategies and targeting two different regions of the *LRRCC1* locus. We could not recover null clones despite repeated attempts in RPE1 - both wild type and p53^-/-^ (Izquierdo et al., 2014), HEK 293 and U2-OS cells, suggesting that a complete lack of LRRCC1 is possibly deleterious. Nevertheless, we obtained partially depleted mutant clones, including three RPE1 clones targeted in either exons 8-9 (clone 1.1) or exons 11-12 (clones 1.2 and 1.9). Clone 1.1 carries deletions in both copies of the *LRRCC1* gene (Supplemental Fig. S3a). However, long in-frame transcripts are expressed at reduced levels through alternative splicing (Supplemental Fig. S1c, S3b). These transcripts are expected to generate mutant protein isoforms carrying deletions in the beginning of the coiled-coil region (Supplemental Fig. S3). In contrast, only wild-type transcripts were detected in clones 1.2 and 1.9, which were present at approximately 30% of wild-type levels, as determined by quantitative RT-PCR (Supplemental Fig. S1c). We could not evaluate the overall decrease in LRRCC1 levels since the endogenous LRRCC1 protein was not detected by Western blot (Supplemental Fig. S1b). However, we confirmed the decrease in centrosomal LRRCC1 levels by immunofluorescence using the two different anti- LRRCC1 antibodies (Fig. 4a; Supplemental Fig. S1d-e). The down-regulation of LRRCC1 in CRISPR clones was overall of the same order as that achieved by RNAi, although treatment of CRISPR clones with the more efficient siRNA (si LRRCC1-1) could further reduce LRRCC1 levels (Fig. 4a). Using Airyscan microscopy, we showed that LRRCC1 amounts were decreased not only at centriolar satellites, but also at the centrioles themselves in CRISPR clones (Fig. 4b). Interestingly, the decrease in centriolar LRRCC1 was less for clone 1.1 than for the other clones, suggesting that the mutant isoforms produced in this clone have different dynamics than wild-type LRRCC1. Following induction of ciliogenesis, the proportion of ciliated cells was decreased in all three mutant clones compared to control cells (Fig. 4c). We were unable to obtain stable RPE1 cell lines expressing tagged versions of LRRCC1, and transient overexpression of LRRCC1 in wild-type cells led to a decrease in the proportion of ciliated cells, making phenotype rescue experiments difficult to interpret. However, we used RNAi as an independent method to verify the specificity of ciliary defects observed in CRISPR clones. The proportion of ciliated cells was decreased by RNAi to a similar extent than in CRISPR clones (Fig. 4c; Supplemental Fig. S1f). RNAi treatment of CRISPR clones did not lead to a greater decrease in ciliary frequency, suggesting that loss of LRRCC1 only partially inhibits ciliogenesis (Fig. 4c). Sensory ciliopathies like JBTS result to a large extent from defective Hedgehog signaling (Romani et al., 2013). We determined the effect of LRRCC1- depletion on Hedgehog signaling by measuring the ciliary accumulation of the activator SMOOTHENED (SMO) upon induction of the pathway (Rohatgi et al., 2007). Depletion of LRRCC1 by either CRISPR editing or RNAi led to a drastic decrease in SMO accumulation at the primary cilium following induction of the Hedgehog pathway by SMO-agonist (SAG) (Fig. 4d, e), and reduced expression of the target gene *PTCH1* (Supplemental Fig. S1i) (Goodrich et al., 1996). Taken together, our results demonstrate that LRRCC1 is required for proper ciliary assembly and signaling in human cells, further establishing its implication in JBTS.

**Figure 4.**
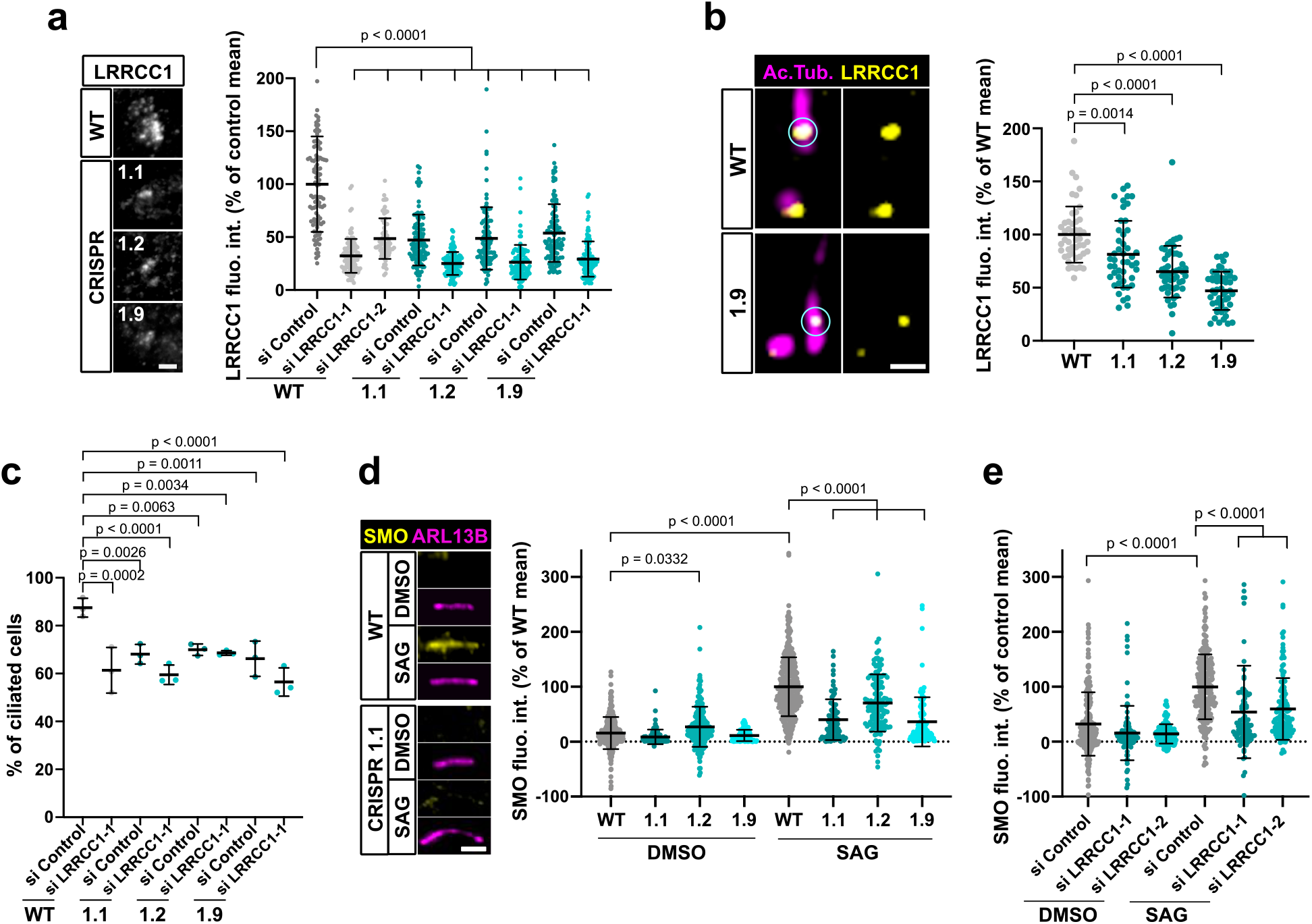
LRRCC1 is required for ciliary assembly and signaling. **a)** Left: LRRCC1 staining (Ab2) of WT or LRRCC1-defficient RPE1 cells obtained by CRISPR/Cas9 editing (clones 1.1, 1.2 and 1.9). Bar, 2 µm. Right: quantification of fluorescence intensity in WT or CRISPR clones treated with control or LRRCC1 siRNAs. Bars, mean ± SD, 3 independent experiments. p values are provided when statistically significant from the corresponding control (One-way ANOVA). **b)** Quantification of LRRCC1 distal pool at the mother centriole of ciliated WT or CRISPR cells. Left: Airyscan images showing the region of interest (circled). LRRCC1 (yellow), acetylated tubulin (magenta). Bar: 0.5 µm. Right: quantification of the corresponding signal. Bars, mean ± SD, ≥ 47 cells from 2 independent experiments. p values are provided when statistically significant from the corresponding control (One-way ANOVA). **c)** Percentage of ciliated cells in WT or CRISPR cells treated with control or LRRCC1 siRNAs and serum-deprived during 24 hours. Bars, mean ± SD, ≥ 204 cells from 3 independent experiments for each condition. p values are provided when statistically significant from the corresponding control (One-way ANOVA). **d)** Left: SMO (yellow) accumulation at primary cilia (ARL13B, magenta) following SAG-induction of the Hedgehog pathway, in WT or CRISPR cells. Bar, 2 µm. Right: quantification of ciliary SMO expressed as a percentage of the SAG-treated WT mean. Bars, mean ± SD, 3 independent experiments. p values are provided when statistically significant from the corresponding control (One-way ANOVA). **e)** Ciliary SMO expressed as a percentage of the SAG-induced control mean in RPE1 cells treated with control or LRRCC1 siRNAs. Bars, mean ± SD, 3 independent experiments. p values are provided when statistically significant from the corresponding control (One-way ANOVA).

### Depletion of LRRCC1 induces defects in centriole structure

Mutations in distal centriole components can alter centriole length regulation or the assembly of DAs, which both result in defective ciliogenesis (Reiter and Leroux, 2017; Sharma et al., 2021). We used U-ExM to search for possible defects in centriole structure in LRRCC1- depleted RPE1 cells. We measured centriole length in CRISPR clone 1.9, which has the lowest levels of centriolar LRRCC1 (Fig. 4b), and in clone 1.1, which expresses mutant isoforms of LRRCC1. Centrioles were co-stained with anti-acetylated tubulin and an antibody against the DA component CEP164 to differentiate mother and daughter centrioles. We observed an increase in centriole length in clone 1.9 (Fig. 5a) compared to control cells (483 ± 53 nm for mother and 372 ± 55 nm for daughter centrioles in clone 1.9; 427 ± 56 nm for mother and 320 ± 46 nm for daughter centrioles in control cells; mean ± SD). Although on a limited sample size, we also observed abnormally long centrioles by transmission electron microscopy in this clone (494 ± 73 nm in clone 1.9, N = 9; 429 ± 52 nm in control cells, N = 3; mean ± SD) (Fig. 5c). The increase in centriole length was not due to mitotic delay as previously observed (Kong et al., 2020), since the duration of mitosis in clone 1.9 was similar as in control cells (Supplemental Fig. S1k). In addition, although centriole length was not modified in clone 1.1, further reduction of LRRCC1 levels by RNAi resulted in a significant increase in centriole length compared to control cells (Fig. 5b). Next, we analyzed DA organization by labeling CEP164, which localizes to the outer part of DAs (Fig. 5d) (Yang et al., 2018). In RPE1 control cells, 80 ± 14 % of mother centriole had 9 properly shaped DAs, but this proportion fell to 57 ± 16 % and 44 ± 17 % (mean ± SD) in clones 1.1 and 1.9, respectively (Fig. 5e). Mutant clones exhibited an increased proportion of centrioles with one or more abnormally shaped DAs (29 ± 17 % and 42 ± 18 % in clones 1.1 and 1.9, respectively, compared to 11 ± 11 % in control cells; mean ± SD). We obtained similar results in a HEK 293 CRISPR clone expressing half the control levels of LRRCC1 (Fig. 5f; Supplemental Fig. S1g). LRRCC1-depletion did not affect overall CEP164 levels at mother centrioles in the CRISPR clones (Supplemental Fig. S4a, d), consistent with the relatively mild defect in DA morphology observed by U-ExM. We also analyzed the distribution of CEP83, a DA component that localizes closer to the centriole wall (Yang et al., 2018). The proportion of centrioles with abnormal CEP83 labeling was not significantly different between control cells and CRISPR clones. However, this proportion became significantly lower than in the control after treating CRISPR clones with RNAi (41 ± 18 % and 48 ± 4 % in clones 1.1 and 1.9 treated with RNAi, respectively, compared to 77 ± 9 % in control cells; mean ± SD; Fig. 5g, h). Beyond these anomalies in centriolar structure, LRRCC1-depleted cells showed no defect in centriole number, supporting that centriole assembly is not affected by LRRCC1 down-regulation (Supplemental Fig. S1j).

**Figure 5.**
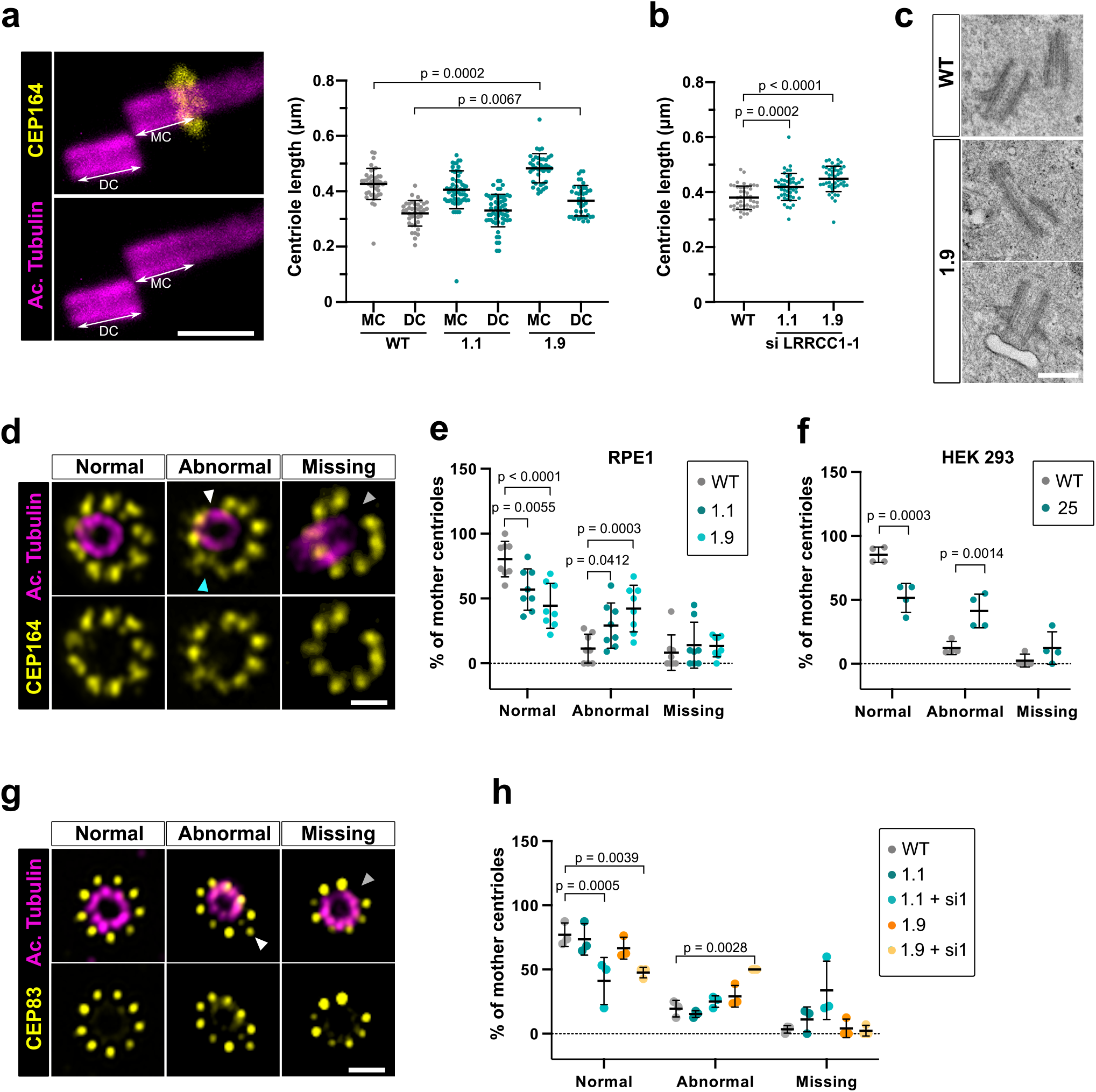
Depleting LRRCC1 induces defects in centriole structure. **a)** Centriole length in mother (MC) and daughter (DC) centrioles analyzed by U-ExM in WT or LRRCC1-deficient clones (1.1 and 1.9). Left: Centrioles were stained for acetylated tubulin (magenta) and CEP164 (yellow) to measure centriole length (arrows). Bar, 0.5 µm. Right: Quantification. Bars, mean ± SD, ≥ 38 centrioles from 3 independent experiments. p values are provided when statistically significant from the corresponding control (One-way ANOVA). **b)** Centriole length in control cells or CRISPR cells treated with LRRCC1 siRNA-1 and stained for acetylated tubulin and CEP83. Bars, mean ± SD, ≥ 43 centrioles from 3 independent experiments. p values are provided when statistically significant from the corresponding control (One-way ANOVA). **c)** Transmission electron microscopy view of centrioles in WT and CRISPR (clone 1.9) RPE1 cells. Note that the 1.9 centrioles are from the same cell. N = 9 centrioles from 8 different cells for clone 1.9, 3 centrioles from 2 different cells for WT. Bar, 0.5 µm. **d)** Examples of normal DAs, DAs with abnormal morphology (white arrowhead: abnormal spacing between consecutive DAs; cyan arrowhead: abnormal DA shape) or missing DAs (grey arrowhead) in RPE1 cells stained with CEP164 (yellow) and analyzed by U-ExM. Images are maximum intensity projections of individual z-sections encompassing the CEP164 signal. Note that an apparent shift between channels occurs when centrioles are slightly angled with respect to the imaging axis. Bar, 1 µm. **e)** Percentages of centrioles presenting anomalies in CEP164 staining in WT or CRISPR RPE1 cells. ≥ 87 centrioles from 8 independent experiments for each condition. p values are provided when statistically significant from the corresponding control (Two-way ANOVA). **f)** Percentages of centrioles presenting anomalies in CEP164 staining in WT or CRISPR HEK 293 (clone 25) cells. ≥ 40 centrioles from 4 independent experiments for each condition. p values are provided when statistically significant from the corresponding control (Two-way ANOVA). **g)** Examples of normal DAs, DAs with abnormal morphology (white arrowhead) or missing DAs (grey arrowhead) in RPE1 cells stained with CEP83 (yellow) and analyzed by U-ExM. Images are maximum intensity projections of individual z- sections encompassing the CEP83 signal. Note that apparent shift between channels and decreased circularity occur when centrioles are slightly angled with respect to the imaging axis. Bar, 1 µm. **h)** Percentages of centrioles presenting anomalies in CEP83 staining in WT RPE1 cells and CRISPR clones with or without RNAi treatment. ≥ 56 centrioles from 3 independent experiments for each condition. p values are provided when statistically significant from the corresponding control (Two-way ANOVA).

Together, these results show that down-regulation of LRRCC1 affects the formation of centriole distal structures, leading to centriole over-elongation and abnormal DA morphology.

### LRRCC1 and C2CD3 delineate a rotationally asymmetric structure in human centrioles

We next wanted to determine whether LRRCC1 cooperates with other distal centriole components. Proteins shown to be recruited early at procentriole distal end include CEP290 (Kim et al., 2008), OFD1 (Singla et al., 2010) and C2CD3 (Thauvin-Robinet et al., 2014). Of particular interest, OFD1 and C2CD3 are required for DA assembly and centriole length control, and mutations in these proteins have been implicated in sensory ciliopathies (Singla et al., 2010; Thauvin-Robinet et al., 2014; Tsai et al., 2019; Lei Wang et al., 2018). We first determined whether depletion of LRRCC1 either by CRISPR editing or by RNAi led to modifications in the recruitment of these proteins within centrioles. We found no major differences in the centrosomal levels of OFD1 and CEP290 compared to control cells (Supplemental Fig. S4b, c, e, f). In contrast, C2CD3 levels were moderately increased in cells depleted from LRRCC1 either by CRISPR editing (clones 1.1 and 1.9) or by RNAi (Fig. 6a, b). We thus analyzed C2CD3 further by U-ExM. As described previously, C2CD3 localized principally at the distal extremity of centrioles (Fig. 6c) (Tsai et al., 2019; Yang et al., 2018). Strikingly, the C2CD3 labeling was also asymmetric, often adopting a C-shape (Fig. 6d). After correcting the vertical alignment of centrioles as previously, we generated an average 3D reconstruction of the C2CD3 pattern. To do this, we used one end of the C as a reference point in the xy-plane to superimpose individual centriole views. The resulting image supported that the C2CD3 labeling forms a C-shaped pattern positioned symmetrically in the centriole lumen (Fig. 6e). To determine whether the C2CD3 localization pattern is affected by LRRCC1- depletion, we next analyzed C2CD3 in LRRCC1 CRISPR clones 1.1 and 1.9. The C2CD3 pattern was more variable than in control RPE1 cells, and often appeared abnormal in shape, position, or both (Fig. 6f). Indeed, averaging the signal from multiple LRRCC1-depleted centrioles produced aberrant patterns, most strikingly for clone 1.9 (clone 1.9; Fig. 6g). Furthermore, the phenotype of clone 1.1 was enhanced by further reducing LRRCC1 levels using RNAi (Fig. 6g). Thus, LRRCC1 is required for the proper assembly of the C2CD3- containing distal structure.

**Figure 6.**
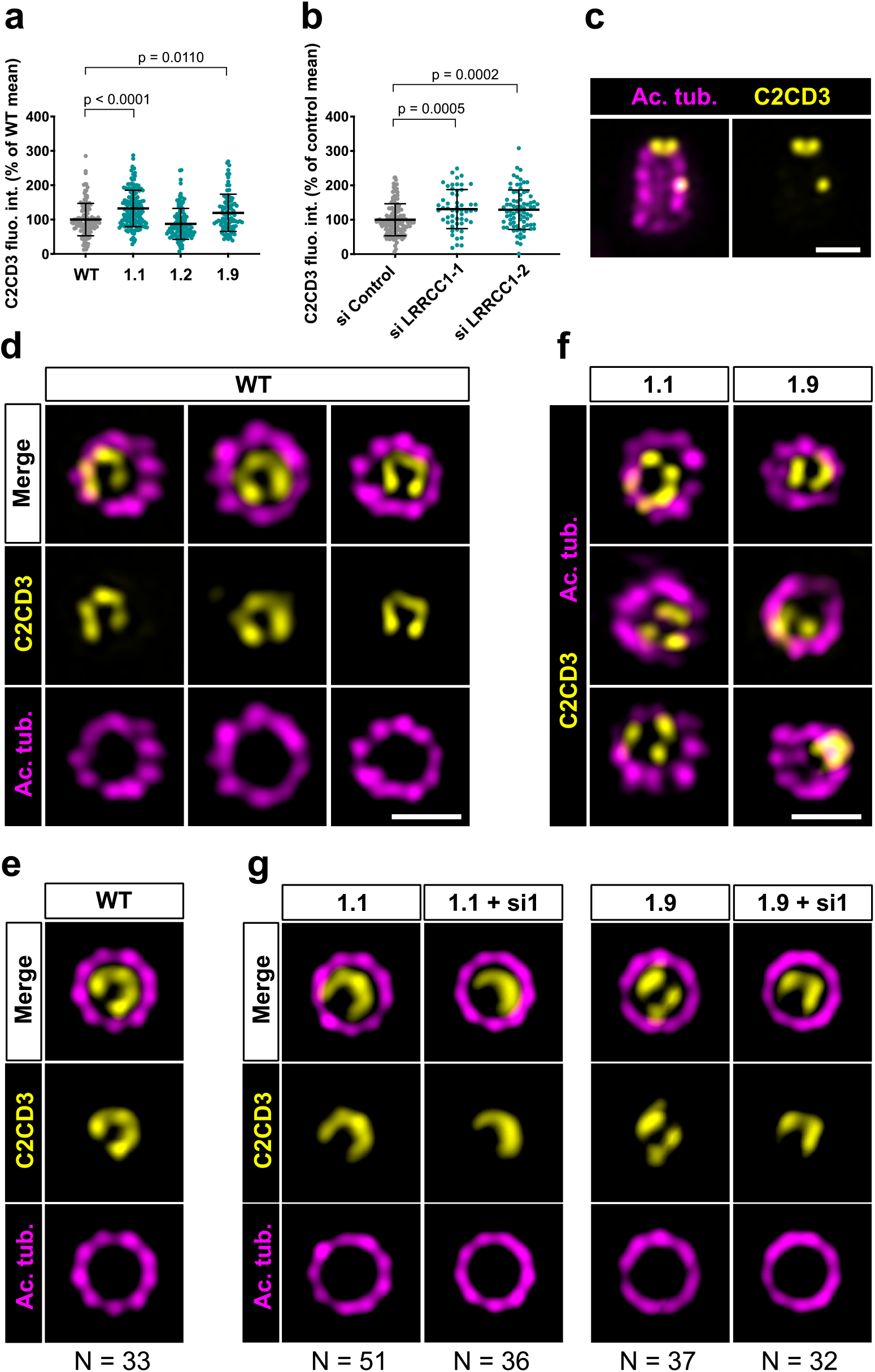
C2CD3 localizes asymmetrically at the distal end of centrioles and is affected by LRRCC1-depletion. **a)** C2CD3 levels at the centrosome of WT or CRISPR RPE1 cells. Bars, mean ± SD, 3 independent experiments. p values are provided when statistically significant from the corresponding control (One-way ANOVA). **b)** C2CD3 levels at the centrosome in RPE1 cells treated with control or LRRCC1 siRNAs. Bars, mean ± SD, 3 independent experiments. p values are provided when statistically significant from the corresponding control (One-way ANOVA). **c)** Longitudinal view of a centriole analyzed by U-ExM and stained for C2CD3 (yellow) and acetylated tubulin (magenta). Bar, 0.2 µm. **d)** Centrioles from WT RPE1 cells as viewed from the distal end. C2CD3 (yellow), acetylated tubulin (magenta). Images are maximum intensity projections of individual z-sections encompassing the C2CD3 signal. Note that an apparent shift between channels occurs when centrioles are slightly angled with respect to the imaging axis. Bar, 0.2 µm. **e)** Average C2CD3 images obtained from 33 individual centrioles from WT RPE1 cells viewed from the distal end, in transverse views. One end of the C-pattern was used as a reference point to align individual centrioles. **f)** Centrioles from untreated CRISPR cells or CRISPR cells treated with LRRCC1 RNAi in transverse section as viewed from the distal end. C2CD3 (yellow), acetylated tubulin (magenta). Images are maximum intensity projections of individual z-sections encompassing the C2CD3 signal. Note that an apparent shift between channels occurs when centrioles are slightly angled with respect to the imaging axis. Bar, 0.2 µm. **g)** Average C2CD3 images obtained from untreated or RNAi-treated CRISPR cells viewed from the distal end, in transverse views. The number or individual centrioles used for generating each average is indicated.

To determine whether LRRCC1 and C2CD3 might belong to a common structure, we next examined their respective positions within the centriole. We co-stained centrioles with our anti- LRRCC1 antibody and a second anti-C2CD3 antibody produced in sheep (Table 1). We confirmed that LRRCC1 and C2CD3 are present in the same distal region of the centriole (Fig. 7a). In transverse views, the two proteins were usually not perfectly colocalized but found in close vicinity of one another near the microtubule wall. However, C2CD3 distal staining was consistently fainter than with the previous antibody, and we either could not observe a full C-shaped pattern, or we could not image it due to fluorescence bleaching. Neither anti-C2CD3 antibodies worked in mouse, so we were not able to compare C2cd3 and Lrrcc1 localization in MCCs. Nevertheless, the results obtained by individually labeling LRRCC1 and C2CD3 at the centrosome (Fig. 1f, 6e) together with the co-localization data (Fig. 7a) are consistent with the hypothesis that LRRCC1 is located along the C2CD3-containing, C-shaped structure (Fig. 7b). C2CD3 was not co-immunoprecipitated with a GFP-LRRCC1 fusion protein however, suggesting that LRRCC1 and C2CD3 do not directly interact (Supplemental Fig. S5).

**Figure 7.**
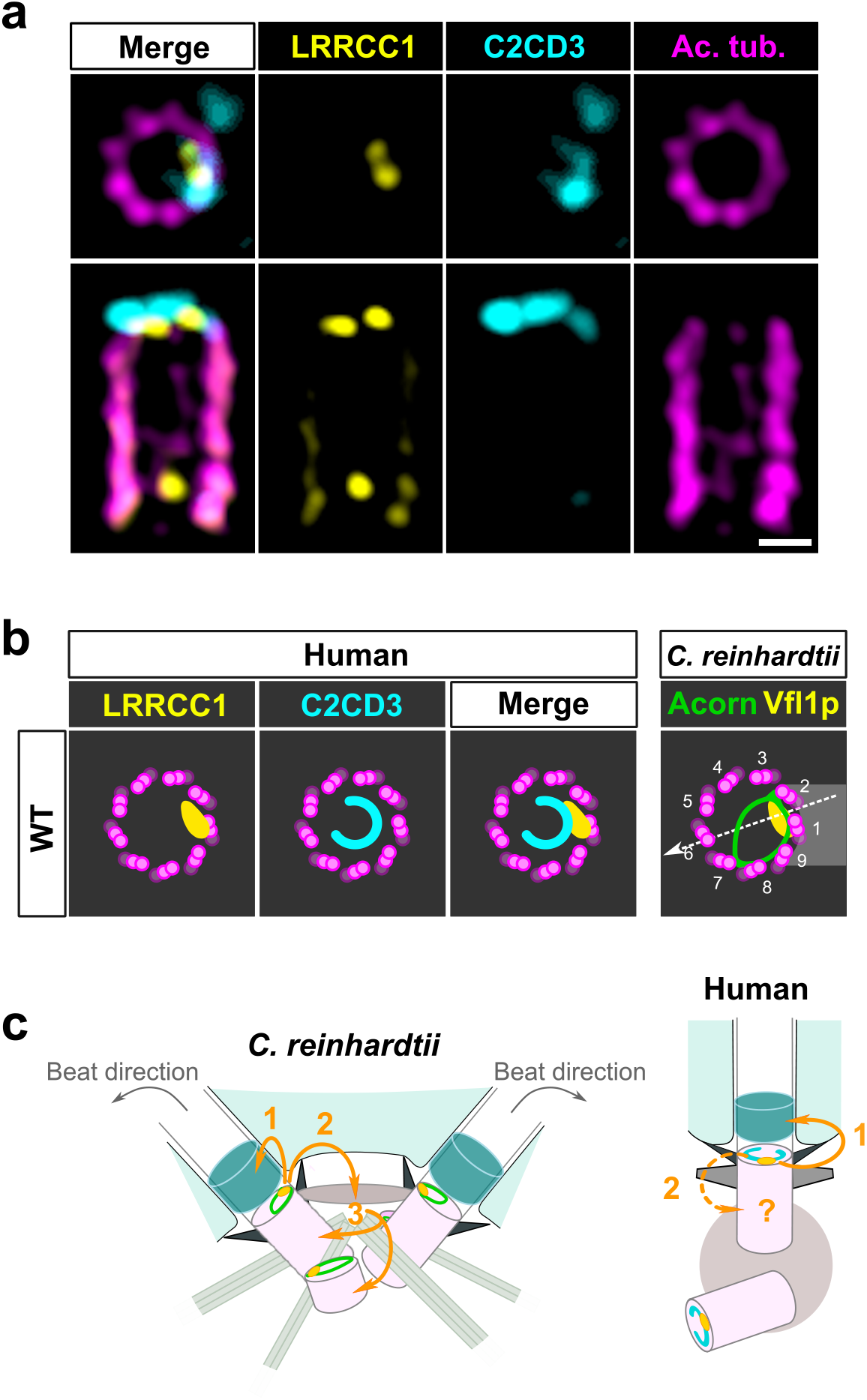
C2CD3 and LRRCC1 partially colocalize at the distal end of centrioles. **a)** RPE1 centrioles processed for U-ExM and stained for LRRCC1 (Ab2, yellow), C2CD3 (cyan) and acetylated tubulin (magenta). Bar, 0.1 µm. **b)** Model showing the possible location of LRRCC1 and C2CD3 relative to each other within human centrioles. Right panel: diagram showing the respective positions of the acorn (Geimer and Melkonian, 2004) and Vfl1p (Silflow et al., 2001) in *C. reinhardtii*. The direction of the flagellar beat is indicated by a dotted arrow, and the distal striated fiber is in grey. **c)** Evolution of the roles played by Vfl1p/LRRCC1 proteins and associated rotationally asymmetric centriolar substructures. In *C. reinhardtii*, Vfl1p is required for proper ciliary assembly (1), as well as for the formation of fibers and microtubular roots (2) that control the position of centrioles and procentrioles (3), and overall cellular organization (Adams et al., 1985; Silflow et al., 2001). In human cells, LRRCC1 and C2CD3 are required for primary cilium assembly (1) - this study and (Thauvin-Robinet et al., 2014; Ye et al., 2014) - and a role in asymmetric anchoring of cytoskeletal elements to the centriole may also be conserved (2), which could indirectly affect the determination of procentriole assembly site.

Taken together, our results support that C2CD3 localizes asymmetrically in the distal lumen of human centrioles, a pattern that depends in part on LRRCC1.

## Discussion

Here, we show that centrioles within the human centrosome are rotationally asymmetric despite the apparent nine-fold symmetry of their ultrastructure. This asymmetry is manifested by a specific enrichment in LRRCC1 near two consecutive triplets, and the C-shaped pattern of C2CD3. Depletion of LRRCC1 perturbed the recruitment of C2CD3 and induced defects in centriole structure, ciliogenesis and ciliary signaling, supporting that LRRCC1 contributes to organizing the distal centriole region together with C2CD3. LRRCC1 localizes like its *C. reinhardtii* ortholog Vfl1p, and C2CD3 delineates a filamentous structure reminiscent of the acorn first described in *C. reinhardtii* (Geimer and Melkonian, 2005, 2004) and later found in a wide variety of eukaryotic species (Cavalier-Smith, 2021; Vaughan and Gull, 2016). Collectively, our results support that rotational asymmetry is a conserved property of centrioles linked to ciliary assembly and signaling in humans.

### LRRCC1 and C2CD3 belong to a conserved rotationally asymmetric structure

Our work identifies two proteins located asymmetrically in the distal centriole lumen of the human centrosome, each with a specific pattern. LRRCC1 localizes principally near two consecutive triplets, with the first triplet counterclockwise labelled approximately 50 % more than the next one. This pattern is highly reminiscent of the LRRCC1 ortholog Vfl1p, which localizes predominantly to the triplet facing the second centriole (referred to as triplet 1), and to a lesser extent to its immediate neighbor counterclockwise (triplet 2; Fig. 7b) (Silflow et al., 2001). In *C. reinhardtii*, triplets 1 and 2 are positioned directly opposite to the direction of flagellar beat, which is directed towards triplet 6 (Fig. 7b) (Lin et al., 2012). In mouse MCCs, Lrrcc1 is associated to triplets located not exactly opposite to the basal foot but with a clockwise shift of at least 20° from the basal foot axis. However, the beating direction was shown to be also shifted approximately 20° clockwise relative to the position of the basal foot in bovine tracheal MCCs (Schneiter et al., 2021) (Fig. 2d). The position of Lrrcc1/Vfl1p-labelled triplets with respect to ciliary beat direction might thus be similar in *C. reinhardtii* and in animal MCCs. Overall, the specific localization pattern of Vfl1p-related proteins at the distal end of centrioles, and their requirement for centriole positioning and ciliary beat orientation when motile cilia are present, appear to be conserved between flagellates and animals.

The second protein conferring rotational asymmetry to human centrioles, C2CD3, delineates a C-shape in the distal lumen. Strikingly, this staining is reminiscent of a filament observed by electron microscopy, which is said to form an ‘incomplete circle’ in the distal lumen of human centrioles (Vorobjev and Chentsov, 1980). Several lines of evidence favor the hypothesis that the C2CD3-containing structure is homologous to the acorn, a conserved filamentous structure that in *C. reinhardtii* connects five consecutive triplets along the centriole wall and across the lumen (Fig. 7b) (Cavalier-Smith, 2021; Geimer and Melkonian, 2004; Vaughan and Gull, 2016). First, the C2CD3 labeling is consistent with a circular filament. Second, C2CD3 is partially co-localized with LRRCC1 near the microtubule wall. And last, C2CD3 orthologs are found in a variety of flagellated unicellular eukaryotes, including the green algae *Micromonas pusilla* (Zhang and Aravind, 2012) and *Chlamydomonas eustigma* (Uniprot_A0A250XH15), suggesting an ancestral association to centrioles and cilia. The partial co-localization of Vfl1p and the acorn in *C. reinhardtii*, and the observation that both are already present at the distal end of procentrioles, led to propose that Vfl1p might also be a component of the acorn (Geimer and Melkonian, 2004). Consistent with this idea, both LRRCC1 and C2CD3 are recruited early to the distal end of human procentrioles, and LRRCC1 is required for proper assembly of the C2CD3-containing structure. C2CD3 recruitment at the centrioles also depends on the proteins CEP120 and Talpid3 (Tsai et al., 2019). Future work will help deciphering the relationships between these different proteins and characterize in more details the architecture of the rotationally asymmetric structure at the distal end of mammalian centrioles.

### Rotationally asymmetric centriole components are required for ciliogenesis

Our results uncover a link between centriole rotational asymmetry and primary ciliogenesis in human cells. Mutations in C2CD3 have been involved in several sensory ciliopathies, including JBTS (Boczek et al., 2018; Cortés et al., 2016; Ooi, 2015; Thauvin-Robinet et al., 2014). The associated ciliary defects are likely caused by anomalies in the structure of centrioles, since depleting C2CD3 inhibits centriole elongation and DA assembly, whereas C2CD3 overexpression leads to centriole hyper-elongation (Thauvin-Robinet et al., 2014; Wang et al., 2018; Ye et al., 2014). We observed similar defects in LRRCC1-depleted cells, but of comparatively lesser extent. DA morphology was altered, and centriole length was slightly increased in cells depleted from LRRCC1. The fact that LRRCC1-depletion has a more limited impact on centriole assembly than perturbation of C2CD3 levels suggests that LRRCC1 might not be directly involved in centriole length control or DA formation, however. The defects observed in LRRCC1-depleted cells could instead result indirectly from the abnormal localization of C2CD3. Besides the defects in centriole structure, it is plausible that LRRCC1- depletion also perturbs the organization of the ciliary gate, as LRRCC1-depleted cells exhibited a drastic reduction in Hedgehog signaling. Loss of ciliary gate integrity interferes with the accumulation of SMO in the cilium upon activation of the Hedgehog pathway and is a frequent consequence of ciliopathic mutations (Garcia-Gonzalo and Reiter, 2017). The ciliary gate consists of the TZ and the DA region, which both contribute to regulating protein trafficking in and out of the cilium (Garcia-Gonzalo and Reiter, 2017; Nachury, 2018). The anomalies in DA morphology observed in LRRCC1-depleted cells could disrupt the organization of the so-called DA matrix (Yang et al., 2018), thus preventing SMO accumulation in the cilium.

Another, non-mutually exclusive possibility is that the architecture of the TZ, which forms directly in contact with the distal end of the centriole, is altered by LRRCC1-depletion. In either case, our observations in RPE1 cells are consistent with the JBTS diagnosis in two siblings carrying a mutation in the *LRRCC1* gene (Shaheen et al., 2016), further establishing that *LRRCC1* is a novel ciliopathy gene. Besides signaling, ciliary gate integrity is required for axoneme extension and indeed, LRRCC1-depleted cells formed cilia at lower frequency than control cells – a defect that might also result from perturbed DA architecture. In the *vfl1* mutant of *C. reinhardtii*, both unanchored centrioles, and centriole docked at the plasma membrane but lacking a flagellum were observed (Adams et al., 1985). This supports that LRRCC1/Vfl1p requirement for properly assembling the ciliary gate is a conserved functional aspect of this family of proteins (Fig. 7c).

Why is there a rotationally asymmetric structure at the base of primary cilia, and how does this structure form and contribute to the assembly of the DAs and the cilium remain open questions. In *C. reinhardtii* and in MCCs, LRRCC1 function is linked to the assembly of asymmetric appendages, which must be correctly positioned in relation to ciliary beat direction (Fig. 7c). An asymmetric structure present early during centriole assembly and ultimately located near the cilium appears well suited for this task. The conservation of such a structure at the base of the primary cilium could perhaps indicate that primary cilia also possess rotationally asymmetric features, which would open interesting new perspectives on ciliary roles in heath and disease.

### Other roles for centriole rotational asymmetry in human cells

Our finding that procentrioles do not form completely at random with respect to LRRCC1 location in the parent centriole suggests that centriole rotational polarity can influence centriole duplication in human cells. In *C. reinhartdtii*, procentrioles are formed at fixed positions with respect to the parent centrioles, to which they are bound by a complex array of fibrous and microtubular roots (Fig. 7c) (Geimer and Melkonian, 2004; Yubuki and Leander, 2013). The process is likely different at the centrosome since the roots typical of flagellates are not conserved in animal cells (Azimzadeh, 2021; Yubuki and Leander, 2013). In mammalian cells, procentrioles form near the wall of the parent centriole following the recruitment of early centriole proteins directly to the PCM components CEP152 and CEP192 (Yamamoto and Kitagawa, 2021). It is nonetheless conceivable that an asymmetry in triplet composition could result in local changes in PCM composition, which in turn could negatively impact PLK4 activation in this region. For instance, our analyses in planarian MCCs led us to postulate that linkers might be tethered to one side of the centrioles in a VFL1-dependent manner and independently of centriole appendages (Basquin et al., 2019). Future work will allow deciphering how centriole rotational asymmetry influences centriole duplication, and whether it affects other aspects of centriole positioning and cellular organization.

## Material and Methods

### Cell culture

RPE1 cells (hTERT-RPE1, RRID:CVCL_4388) were cultured in DMEM/F-12 medium (ThermoFisher Scientific) supplemented with 10 % fetal calf serum (ThermoFisher Scientific), 100 U/mL penicillin and 100 μg/mL streptomycin (ThermoFisher Scientific). Ciliogenesis was induced by culturing RPE1 cells in medium without serum during 48 hours. HEK 293 cells (kind gift from F. Causeret, Institut Imagine, Paris) were cultured in DMEM medium (ThermoFisher Scientific) supplemented with 10 % fetal calf serum and antibiotics as previously. All cells were kept at 37°C in the presence of 5 % CO2.

### Mouse ependymal cells and tracheal tissue

All experiments were performed in accordance with French Agricultural Ministry and European guidelines for the care and use of laboratory animals. *In vitro* differentiated ependymal cells were a kind gift from A.R. Boudjema and A. Meunier (IBENS, Paris). They were prepared as described previously (Delgehyr et al., 2015; Mercey et al., 2019) from Cen2GFP mice (CB6-Tg(CAG-EGFP/CETN2)3-4Jgg/J, The Jackson Laboratory). The fragment of trachea was obtained from a wild-type mouse of the Swiss background (kindly provided by I. Le Parco, Institut Jacques Monod).

### CRISPR/Cas9 editing

LRRCC1 mutant clones were obtained by two different CRISPR/Cas9 strategies. First, RPE1 cells were co-transfected with plasmid px154-1 (U6p-gRNA#1_U6p-gRNA#2_CMVpnCas9- EGFP_SV40p-PuroR-pA with gRNA#1: 5’- AGA ATT CTA CCC TAC CTG - 3’ and gRNA#2: 5’- TAA GGT AGT GCT TCC TAC -3’) targeting the *LRRCC1* locus in exon 8, and px155-24 (U6p-gRNA#3_U6p-gRNA#4_CMVpnCas9-mCherry_SV40p-PuroR-pA; gRNA#1: 5’- ATC TAC TCG GAA AGC TGA -3’ and 5’- GCT TGA GGG CTC AAA TAC- 3’) targeting exon 9. Both constructs express the nickase mutant of Cas9 fused to either EGFP or mCherry. Two days after transfection, EGFP- and mCherry-positive cells were sorted by flow cytometry and grown at low concentration. Individual clones were picked after 2 weeks and analyzed by PCR to detect short insertions/deletions. A single clone was obtained (clone 1.1), which was further characterized by genomic sequencing. Both alleles of *LRRCC1* contained deletions (∼ 0.6 kb deletion of exon 9 and a ∼1.5 kb deletion of exon 8; Supplemental Fig. S3a) leading to frameshifts. In a second approach, cells were co-transfected using a mix of 3 CRISPR/Cas9 Knockout Plasmids (sc-413781; Santa Cruz Biotechnology) targeting exons 11 (5’- CTT GTT CTC TTT CTC GAT GA– 3’ and 5’ - ACT TCT TGC ATT GAA AGA AC- 3’) or 12 (5’ - CGT GTT AAG CCA GCA GTA TA– 3’) of *LRRCC1,* together with the corresponding Homology Directed Repair plasmids carrying a puromycine-resistance cassette (sc-413781-HDR; Santa Cruz Biotechnology), following the recommendations of the manufacturer. Mutant clones were selected by addition of 2 µg/mL puromycine in the culture medium and further screened by immunofluorescence, allowing to identify two clones with decreased LRRCC1 levels (clones 1.2 and 1.9). Genomic insertion of the HDR cassette could not be detected in these clones by PCR, and no sequence anomalies were identified in PCR fragments corresponding to exons 10 to 13. This suggests that one copy of the *LRRCC1* gene is intact, while the second copy may have undergone more extensive modifications via large deletions/insertions. For sequencing of LRRCC1 transcripts, total RNA extracts were obtained using the Nucleospin RNA kit (Macherey-Nagel) and cDNAs were synthesized using SuperScript III reverse transcriptase (Thermofisher Scientific). PCR primers specific to exons 4 and 8, 4 and 9, 8 and 19, or 9 and 19 were used to amplify cDNAs from clone 1.1; primers specific to exons 4 and 17 were used for clones 1.2 and 1.9. The resulting fragments were analyzed by sequencing.

### Inducible HEK 293 cell lines

LRRCC1 full-length coding sequence was amplified from cDNA clone IMAGE:5272572 (Genbank accession: BC070092.1), corresponding to the longest isoform of LRRCC1 (NM_033402.5), after correction of a frameshift error by PCR mutagenesis. As N- and C-terminal GFP fusions were not targeted to the centrosome, we inserted the GFP tag within the LRRCC1 sequence in disordered regions present between the leucine rich repeat and coiled-coil domains, either after amino acid 251 or 402. The fusions were cloned into the pCDNA- 5FRT (ThermoFischer Scientific) vector using the Gibson assembly method (Gibson et al., 2009) and then integrated into the Flp-In-293 cell line using the Flp-In system (ThermoFischer Scientific). Expression of the GFP-LRRCC1 fusions was induced by culturing the Flp-In-293 cell lines overnight in medium supplemented with 1 µg/mL doxycycline (ThermoFischer Scientific).

### RNAi

Ready to use double-stranded siRNA LRRCC1-si1 (target sequence: 5’- AAG GAG AAA GAT GGA GAC GAT - 3’) (Muto et al., 2008), LRRCC1-si2 (target sequence: 5’- TTA GAT GAC CAA ATT CTA CAA - 3’) and control siRNA (AllStars Negative Control) were purchased from Qiagen. siRNAs were delivered into cells using Lipofectamine RNAiMAX diluted in OptiMEM medium (ThermoFisher Scientific). Cells were fixed after 48 hours and processed for immunofluorescence. For RNAi-depletion of ciliated cells, RPE1 cells grown in complete culture medium were treated by RNAi, incubated for 2 days, then submitted to a second round of RNAi. After 8 hours, cells were washed 3 x in PBS then cultured during 24 hours in serum-free medium to induce ciliogenesis.

### qRT-PCR

Total RNA extracts were obtained using the Nucleospin RNA kit (Macherey-Nagel) and cDNAs were synthetized using SuperScript III reverse transcriptase (Thermofisher Scientific). qPCR was performed in triplicate with the GoTaq qPCR Master Mix (Promega) in a LightCycler 480 instrument (Roche) using the primers listed in Table 2. Quantification of relative mRNA levels was performed using CHMP2A and EMC7 as reference genes following the MIQE guidelines (Bustin et al., 2009).

### Antibodies

Fragments encoding either aa 671-805 (Ab1) or aa 961-1032 (Ab2) of LRRCC1 (NP_208325.3) were cloned in pGST-Parallel1 and expressed in *Escherichia coli.* The GST-fusion proteins were purified under native conditions using glutathione agarose (ThermoFisher Scientific) and the LRRCC1 fragments were recovered by Tev protease cleavage and dialyzed before rabbit immunization (Covalab). Antibodies were affinity-purified over the corresponding GST-LRRCC1 fusion bound to Affi-Gel 10 resin (Bio-Rad). Other primary and secondary antibodies used in this study are listed in Table 1.

### Western blot

For whole-cell extracts, Flp-In-293 cell lines expressing the GFP-LRRCC1 fusions were induced overnight with doxycycline, collected by centrifugation, and resuspended in Western blot sample buffer prior to incubation at 95 °C for 5 minutes. For immunoprecipitation experiments, doxycycline-induced cells expressing LRRCC1 with a GFP inserted after aa 402 were resuspended in lysis buffer (50 mM Tris pH 8, 150 mM NaCl, 1 % NP-40, 0.5 % sodium deoxycholate, 0.1 % SDS) supplemented with 1 mM MgCl2, 20 µg/mL DNAse I (Roche) and a protease inhibitor cocktail (Complete mini, EDTA-free, Roche). After 30 minutes on ice, the lysates were centrifuged at 15 000 g for 10 minutes at 4°C. The supernatants were then incubated with Dynabeads M-280 sheep-anti rabbit magnetic beads (ThermoFischer Scientific) previously incubated with rabbit anti-IgGs, either anti-GFP or anti-HA tag for the control IP (Table 1), and rotated for 3 hours at 4°C. After 3 washes with lysis buffer, immunoprecipitated proteins were recovered by resuspending the beads in sample buffer and heating at 95 °C for 5 minutes. The samples were then run on 4-20% Mini-Protean TGX precast protein gels (Bio-Rad) and transferred onto PVDF membrane using the iBlot 2 blot system (ThermoFischer Scientific). The membranes were blocked and incubated with antibodies following standard procedures, then visualized using Pierce ECL plus chemiluminescence reagents (ThermoFischer Scientific) on a ChemiDoc imaging system (Bio-Rad).

### Immunofluorescence

Cells were fixed in cold methanol for 5 minutes at – 20 °C, blocked 10 minutes with 3 % BSA (Sigma Aldrich) in PBS containing 0.05 % Tween-20 (PBST-0.05%), then incubated with primary antibodies diluted in PBST-0.05% containing 3 % BSA for 1 hour. After washing 3 x 1 minute in PBST-0.05%, cells were incubated 2 hours with secondary antibodies in PBST- 0.05% containing 3 % BSA and 5 µg/mL Hoechst 33342 (ThermoFischer Scientific), washed in PBST-0.05% as previously, and mounted using Fluorescence Mounting Medium (Agilent). For staining of primary cilia with anti-acetylated tubulin, cells were incubated 2 hours on ice prior to methanol fixation. For quantification of SMO accumulation within cilia, confluent cells cultured during 24 hours in serum-free medium were supplemented with 200 nM SAG (Sigma) diluted in DMSO, or DMSO alone for 24 hours. Cells were then co-stained for SMO and ARL13B to determine the position of the primary cilium. For all experiments involving induction of ciliogenesis by serum deprivation, we verified that cells were arrested in G0 by immunofluorescence staining of Ki67. To visualize centriolar LRRCC1, and to quantify CEP290 centrosomal levels, cells were treated during 1 hour with 5 µM nocodazole prior to fixation. Images were acquired using an Axio Observer Z.1 microscope (Zeiss) equipped with a sCMOS Orca Flash4 LT camera (Hamamatsu) and a 63x objective (Plan Apo, N.A. 1.4). The structured illumination microscopy (SIM) image was acquired on an ELYRA PS.1 (Zeiss) equipped with an EMCCD iXon 885 camera (Andor) and a 63x objective (Plan Apo, N.A. 1.4).

### Ultrastructure expansion microscopy

We used the U-ExM protocol described in (Gambarotto et al., 2019) with slight modifications. Cells grown on glass coverslips were incubated in a fresh solution of 1 % acrylamide and 0.7 % formaldehyde diluted in PBS. After incubating 5 hours to overnight at 37 °C, the coverslips were washed with PBS and placed cells down on a drop of 35 μL monomer solution (19.3 % sodium acrylate, 10 % acrylamide, 0.1 % bis-acrylamide in PBS) to which 0.5 % TEMED and 0.1 % ammonium persulfate were added just before use. The coverslips were incubated 5 minutes on ice then 1 hour at 37°C, then transferred to denaturation buffer (200 mM SDS, 200 mM NaCl, 50 mM Tris pH9) for 15 minutes with agitation to detach the gels from the coverslips. The gels were then incubated in denaturation buffer 1.5 hours at 95 °C, washed 2 x 30 minutes in deionized water then incubated overnight in water at room temperature to allow expansion of the gel. The gels were measured at this step to determine the coefficient of expansion. After 2 x 10 minutes in PBS, the gels were cut into smaller pieces then incubated 3 hours at 37 °C with primary antibodies diluted in saturation buffer (3 % BSA, 0.05 % Tween- 20 in PBS). The gel fragments were then washed 3 x 10 minutes in PBST-0.1%, incubated 3 h with secondary antibodies and washed in PBST-0.1% as previously. Finally, the gels were incubated 2 x 30 minutes in de-ionized water then left to expand overnight in de-ionized water to regain their maximum size. For U-ExM of mouse tracheal cells, a fragment of WT mouse trachea (kind gift from I. Le Parco, IJM, Paris) was adhered on a poly-lysine coated coverslip then processed as described above with the following modifications: for the first step, the fragment of trachea was incubated overnight to 48 hours in 1 % acrylamide and 0.7 % formaldehyde in PBS; they were placed 15 min on ice prior to the 1-hour incubation at 37 °C and the transfer to denaturation buffer. Note that GFP fluorescence was quenched during U-ExM processing, so the GFP-Cen2 construct expressed in ependymal cells was not detectable in final samples. Gels were imaged on Lab-Tek chamber slides (0.15 mm) coated with poly-lysine (ThermoFisher Scientific). Images were acquired at room temperature using either a LSM780 confocal microscope (Zeiss) equipped with an oil 63x objective (Plan Apo, N.A. 1.4), or an LSM980 confocal microscope with Airyscan 2 (Zeiss) equipped with an oil 63x objective (Plan Apo, N.A. 1.4).

### Image analysis

Protein levels were determined using ImageJ software (Schneider et al., 2012) by measuring the fluorescence intensity in the centrosome or cilium area and subtracting the cytoplasmic background in z-series taken at 0.5-μm interval. Images of individual centrioles in U-ExM are maximum intensity projections of all z-sections comprising the signal of interest. Note that centrioles are presented as they are in the sample (*i.e.,* without correcting their orientation), which leads to an apparent shift between channels or decreased circularity in the projections when centrioles are not parallel to the imaging axis. Analysis of distal appendage morphology defects was performed on z-stacks and not on projected images. Daughter centriole length was determined by U-ExM using the acetylated tubulin staining. For mother centrioles, which could be associated with a primary cilium, the length was measured between the proximal end of the acetylated tubulin staining and distal appendages labeled by anti-CEP164 or CEP83. To generate average images of LRRCC1 and C2CD3, only centrioles that were nearly perpendicular to the imaging plane were acquired on the Airyscan microscope in order to maximize the resolution in transverse views. Calculating the average image consisted of several steps: cropping out individual centrioles, aligning them, providing reference points, standardizing centrioles using the reference points, and averaging (Supplemental Fig. S2). The cropping was done in ImageJ, and for aligning and providing the reference points a graphical user interface was developed based on Napari (Sofroniew et al., 2020). Centriole alignment: the direction of centriole long axis was selected manually and used to position the centriole vertically. Providing the reference points: reference points were manually selected to outline the circle of microtubules triplets and the location of the protein of interest. The centriole was also framed in Z dimension with a rectangle. Standardization: the reference points were used to calculate all necessary transformations (rotation, scaling and translation) to map the original image of a centriole to the standard image. Averaging: an average image was calculated for all the successive XY planes of the standardized image stacks. For alignment of tracheal cell centrioles, since the current version of the graphical user interface can only accommodate two channels, the position of the basal foot provided by the γ-tubulin channel was reported manually in the acetylated tubulin channel using Image J. The images were then processed as before using the manual annotation as a reference point for the basal foot.

For analysis of procentriole position and LRRCC1 location in procentrioles, 3D- reconstructions of diplosomes processed for U-ExM were obtained using IMARIS software (Oxford Instruments).

### Electron microscopy

RPE1 cells were grown at confluence before induction of ciliogenesis for 72 hours by serum deprivation. Cells were fixed 30 minutes in 2.5 % glutaraldehyde (Electron Microscopy Sciences), 2 % paraformaldehyde (Electron Microscopy Sciences), 1 mM CaCl2 in PBS, then washed 3 x 5 minutes in PBS. Samples were then post-fixed during 30 minutes in 1 % Osmium tetroxide (Electron Microscopy Sciences), then washed 3 x 5 minutes in water. Dehydration was performed using graded series of ethanol in water for 5 minutes 30%, 50%, 70%, 90%, 100%, 100%. Resin infiltration was performed by incubating 30 minutes in an Agar low viscosity resin (Agar Scientific Ltd) and EtOH (1:2) mix, then 30 minutes in a resin and EtOH (2:1) mix followed by overnight incubation in pure resin. The resin was then changed and the samples further incubated during 1.5 hours prior to inclusion in gelatin capsules and overnight polymerization at 60 °C. 70 nm sections were obtained using an EM UC6 ultramicrotome (Leica), post-stained in 4 % aqueous uranyl acetate and lead citrate, and observed at 80 kV with a Tecnai12 transmission electron microscope (ThermoFischer Scientific) equipped with a 1K×1K Keen View camera (OSIS).

### Videomicroscopy

To determine the duration of mitosis, individual frames of cells growing under normal culture conditions were acquired every 5 minutes for 24 hours using an IncuCyte ZOOM live-cell analysis system (Sartorius) equipped with a 20 x objective.

### Statistical analysis

All statistical analyses were performed using the Prism 9 for Mac OS X software (GraphPad Software, Inc.). All values are provided as mean ± SD. The number of experimental replicates and the statistical test used are indicated in the figure legends, and the p values are included when statistically different.

## Acknowledgements

The authors are deeply grateful to Marine Laporte, Virginie Hamel, Paul Guichard and Davide Gambarotto for teaching them the U-ExM procedure and for sharing antibodies; Arnaud Echard and Takashi Ochi for critical reading of the manuscript; Amélie-Rose Boudjema and Alice Meunier for providing mouse ependymal cells and Isabelle Le Parco for the tracheal tissue; Juliane Da Graça and Simon Herman for technical help; Rémi Le Borgne for help with transmission electron microscopy and for critical reading of the manuscript. We acknowledge the core imaging facility of Institut Jacques Monod (ImagoSeine facility, member of the France BioImaging infrastructure supported by grant ANR-10-INBS-04 from the French National Research Agency). This work was supported by funding from La Ligue Contre le Cancer, Fondation ARC pour la recherche sur le cancer and ANR-21-CE13-008 to J.A. N.G. was recipient of a MESRI PhD fellowship from the French Government and a 4^th^ year PhD fellowship from the Fondation pour la Recherche Médicale.

## Supplemental material

Fig. S1 provides additional data on LRRCC1 expression in CRISPR clones and RNAi-treated cells. **Fig. S2** presents the image analysis pipeline for generating the average images of LRRCC1 and C2CD3 staining. **Fig. S3** shows the genomic deletions in the CRISPR clone 1.1 and the corresponding transcripts. **Fig. S4** shows the quantification of the DA component CEP164, and the distal centriole components CEP290 and OFD1, at the centrosome of RPE1 cells depleted from LRRCC1 by CRISPR or RNAi. **Fig. S5** shows that C2CD3 is not co- immunoprecipitated with GFP-LRRCC1.

## References

Adams GM, Wright RL, Jarvik JW. 1985. Defective temporal and spatial control of flagellar assembly in a mutant of Chlamydomonas reinhardtii with variable flagellar number. J Cell Biol 100:955–964.

Andersen JS, Wilkinson CJ, Mayor T, Mortensen P, Nigg EA, Mann M. 2003. Proteomic characterization of the human centrosome by protein correlation profiling. Nature 426:570–574.

Azimzadeh J. 2021. Evolution of the centrosome, from the periphery to the center. Current Opinion in Structural Biology 66:96–103. doi:10.1016/j.sbi.2020.10.020

Azimzadeh J, Hergert P, Delouvée A, Euteneuer U, Formstecher E, Khodjakov A, Bornens M. 2009. hPOC5 is a centrin-binding protein required for assembly of full-length centrioles. Journal of Cell Biology 185. doi:10.1083/jcb.200808082

Basquin C, Ershov D, Gaudin N, Vu HT, Louis B, Papon JF, Orfila AM, Mansour S, Rink JC, Azimzadeh J. 2019. Emergence of a Bilaterally Symmetric Pattern from Chiral Components in the Planarian Epidermis. Dev Cell 51:516–525 e5. doi:10.1016/j.devcel.2019.10.021

Boczek NJ, Hopp K, Benoit L, Kraft D, Cousin MA, Blackburn PR, Madsen CD, Oliver GR, Nair AA, Na J, Bianchi DW, Beek G, Harris PC, Pichurin P, Klee EW. 2018. Characterization of three ciliopathy pedigrees expands the phenotype associated with biallelic C2CD3 variants. European Journal of Human Genetics 26:1797–1809. doi:10.1038/s41431-018-0222-3

Bustin SA, Benes V, Garson JA, Hellemans J, Huggett J, Kubista M, Mueller R, Nolan T, Pfaffl MW, Shipley GL, Vandesompele J, Wittwer CT. 2009. The MIQE guidelines: minimum information for publication of quantitative real-time PCR experiments. Clin Chem 55:611–622. doi:10.1373/clinchem.2008.112797

Cavalier-Smith T. 2021. Ciliary transition zone evolution and the root of the eukaryote tree: implications for opisthokont origin and classification of kingdoms Protozoa, Plantae, and Fungi. Protoplasma. doi:10.1007/S00709-021-01665-7

Clare DK, Magescas J, Piolot T, Dumoux M, Vesque C, Pichard E, Dang T, Duvauchelle B, Poirier F, Delacour D. 2014. Basal foot MTOC organizes pillar MTs required for coordination of beating cilia. Nature communications 5:4888. doi:10.1038/ncomms5888

Cortés CR, McInerney-Leo AM, Vogel I, Rondón Galeano MC, Leo PJ, Harris JE, Anderson LK, Keith PA, Brown MA, Ramsing M, Duncan EL, Zankl A, Wicking C. 2016 Mutations in human C2CD3 cause skeletal dysplasia and provide new insights into phenotypic and cellular consequences of altered C2CD3 function. Scientific Reports 6. doi:10.1038/srep24083

Delgehyr N, Meunier A, Faucourt M, Grau MB, Strehl L, Janke C, Spassky N. 2015. Ependymal cell differentiation, from monociliated to multiciliated cells. Methods in Cell Biology 127:19–35. doi:10.1016/bs.mcb.2015.01.004

Feldman JL, Geimer S, Marshall WF. 2007. The mother centriole plays an instructive role in defining cell geometry. PLoS Biol 5:e149.

Gambarotto D, Zwettler FU, le Guennec M, Schmidt-Cernohorska M, Fortun D, Borgers S, Heine J, Schloetel JG, Reuss M, Unser M, Boyden ES, Sauer M, Hamel V, Guichard P. 2019. Imaging cellular ultrastructures using expansion microscopy (U-ExM). Nature Methods 16:71–74. doi:10.1038/s41592-018-0238-1

Garcia-Gonzalo FR, Reiter JF. 2017. Open Sesame: How Transition Fibers and the Transition Zone Control Ciliary Composition. Cold Spring Harb Perspect Biol 9. doi:10.1101/cshperspect.a028134

Geimer S, Melkonian M. 2005. Centrin scaffold in Chlamydomonas reinhardtii revealed by immunoelectron microscopy. Eukaryot Cell 4:1253–1263.

Geimer S, Melkonian M. 2004. The ultrastructure of the Chlamydomonas reinhardtii basal apparatus: identification of an early marker of radial asymmetry inherent in the basal body. J Cell Sci 117:2663–2674.

Gibson DG, Young L, Chuang RY, Venter JC, Hutchison CA, Smith HO. 2009. Enzymatic assembly of DNA molecules up to several hundred kilobases. Nature Methods 6:343–345. doi:10.1038/nmeth.1318

Goodrich L v., Johnson RL, Milenkovic L, McMahon JA, Scott MP. 1996. Conservation of the hedgehog/patched signaling pathway from flies to mice: Induction of a mouse patched gene by Hedgehog. Genes and Development 10:301–312. doi:10.1101/gad.10.3.301

Gupta GD, Coyaud É, Gonçalves J, Mojarad BA, Liu Y, Wu Q, Gheiratmand L, Comartin D, Tkach JM, Cheung SWT, Bashkurov M, Hasegan M, Knight JD, Lin ZY, Schueler M, Hildebrandt F, Moffat J, Gingras AC, Raught B, Pelletier L. 2015. A Dynamic Protein Interaction Landscape of the Human Centrosome-Cilium Interface. Cell 163:1484–1499. doi:10.1016/j.cell.2015.10.065

Izquierdo D, Wang WJ, Uryu K, Tsou MF. 2014. Stabilization of cartwheel-less centrioles for duplication requires CEP295-mediated centriole-to-centrosome conversion. Cell Rep 8:957–965. doi:10.1016/j.celrep.2014.07.022

Kim J, Krishnaswami SR, Gleeson JG. 2008. CEP290 interacts with the centriolar satellite component PCM-1 and is required for Rab8 localization to the primary cilium. Hum Mol Genet 17:3796–3805. doi:10.1093/hmg/ddn277

Kong D, Sahabandu N, Sullenberger C, Vásquez-Limeta A, Luvsanjav D, Lukasik K, Loncarek J. 2020. Prolonged mitosis results in structurally aberrant and over-elongated centrioles. The Journal of cell biology 219. doi:10.1083/JCB.201910019

Kumar D, Reiter J. 2021. How the centriole builds its cilium: of mothers, daughters, and the acquisition of appendages. Current Opinion in Structural Biology. doi:10.1016/j.sbi.2020.09.006

Le Guennec M, Klena N, Gambarotto D, Laporte MH, Tassin A-M, van den Hoek H, Erdmann PS, Schaffer M, Kovacik L, Borgers S, Goldie KN, Stahlberg H, Bornens M, Azimzadeh J, Engel BD, Hamel V, Guichard P. 2020. A helical inner scaffold provides a structural basis for centriole cohesion. Science Advances 6. doi:10.1126/sciadv.aaz4137

Le Guennec M, Klena N, Aeschlimann G, Hamel V, Guichard P. 2021. Overview of the centriole architecture. Current Opinion in Structural Biology 66:58–65. doi:10.1016/j.sbi.2020.09.015

Lin J, Heuser T, Song K, Fu X, Nicastro D. 2012. One of the Nine Doublet Microtubules of Eukaryotic Flagella Exhibits Unique and Partially Conserved Structures. PLoS ONE 7. doi:10.1371/journal.pone.0046494

Mercey O, al Jord A, Rostaing P, Mahuzier A, Fortoul A, Boudjema AR, Faucourt M, Spassky N, Meunier A. 2019. Dynamics of centriole amplification in centrosome-depleted brain multiciliated progenitors. Scientific Reports 9:1–11. doi:10.1038/s41598-019-49416-2

Meunier A, Azimzadeh J. 2016. Multiciliated cells in animals. Cold Spring Harbor Perspectives in Biology 8. doi:10.1101/cshperspect.a028233

Muto Y, Yoshioka T, Kimura M, Matsunami M, Saya H, Okano Y. 2008. An evolutionarily conserved leucine-rich repeat protein CLERC is a centrosomal protein required for spindle pole integrity. Cell Cycle 7:2738–2748.

Nachury M v. 2018. The molecular machines that traffic signaling receptors into and out of cilia. Current Opinion in Cell Biology. doi:10.1016/j.ceb.2018.03.004

Nommick A, Boutin C, Rosnet O, Schirmer C, Bazellières E, Thomé V, Loiseau E, Viallat A, Kodjabachian L. 2022. Lrrcc1 and Ccdc61 are conserved effectors of multiciliated cell function. Journal of cell science. doi:10.1242/JCS.258960

Ooi J. 2015. Mutations in C2CD3 cause oral-facial-digital syndrome through deregulation of centriole length. Clinical Genetics 87:328–329. doi:10.1111/cge.12545

Reiter JF, Leroux MR. 2017. Genes and molecular pathways underpinning ciliopathies. Nat Rev Mol Cell Biol 18:533–547. doi:10.1038/nrm.2017.60

Rohatgi R, Milenkovic L, Scott MP. 2007. Patched1 regulates hedgehog signaling at the primary cilium. Science 317:372–376. doi:10.1126/science.1139740

Romani M, Micalizzi A, Valente EM. 2013. Joubert syndrome: congenital cerebellar ataxia with the molar tooth. Lancet neurology 12:894–905. doi:10.1016/S1474-4422(13)70136-4

Schneiter M, Halm S, Odriozola A, Mogel H, Rička J, Stoffel MH, Zuber B, Frenz M, Tschanz SA. 2021. Multi-scale alignment of respiratory cilia and its relation to mucociliary function. Journal of Structural Biology 213:107680. doi:10.1016/j.jsb.2020.107680

Shaheen R, Szymanska K, Basu B, Patel N, Ewida N, Faqeih E, al Hashem A, Derar N, Alsharif H, Aldahmesh MA, Alazami AM, Hashem M, Ibrahim N, Abdulwahab FM, Sonbul R, Alkuraya H, Alnemer M, al Tala S, Al-Husain M, Morsy H, Seidahmed MZ, Meriki N, Al-Owain M, AlShahwan S, Tabarki B, Salih MA, Ciliopathy W, Faquih T, El-Kalioby M, Ueffing M, Boldt K, Logan C v, Parry DA, al Tassan N, Monies D, Megarbane A, Abouelhoda M, Halees A, Johnson CA, Alkuraya FS. 2016. Characterizing the morbid genome of ciliopathies. Genome biology 17:242. doi:10.1186/s13059-016-1099-5

Sharma A, Olieric N, Steinmetz MO. 2021. Centriole length control. Current Opinion in Structural Biology 66:89–95. doi:10.1016/j.sbi.2020.10.011

Silflow CD, LaVoie M, Tam LW, Tousey S, Sanders M, Wu W, Borodovsky M, Lefebvre PA. 2001. The Vfl1 Protein in Chlamydomonas localizes in a rotationally asymmetric pattern at the distal ends of the basal bodies. J Cell Biol 153:63–74.

Singla V, Romaguera-Ros M, Garcia-Verdugo JM, Reiter JF. 2010. Ofd1, a human disease gene, regulates the length and distal structure of centrioles. Dev Cell 18:410–424.

Sofroniew N, Lambert T, Evans K, Nunez-Iglesias J, Yamauchi K, Solak AC, Bokota G, ziyangczi, Buckley G, Winston P, Tung T, Pop DD, Hector, Freeman J, Bussonnier M, Boone P, Royer L, Har-Gil H, Axelrod S, Rokem A, Bryant, Kiggins J, Huang M, Vemuri P, Dunham R, Manz T, jakirkham, Wood C, Siqueira A de, Chopra B. 2020. napari/napari: 0.3.8rc1. doi:10.5281/ZENODO.4046812

Takao D, Yamamoto S, Kitagawa D. 2019. A theory of centriole duplication based on self- organized spatial pattern formation. Journal of Cell Biology 218:3537–3547. doi:10.1083/JCB.201904156

Thauvin-Robinet C, Lee JS, Lopez E, Herranz-Pérez V, Shida T, Franco B, Jego L, Ye F, Pasquier L, Loget P, Gigot N, Aral B, Lopes CAM, St-Onge J, Bruel AL, Thevenon J, González-Granero S, Alby C, Munnich A, Vekemans M, Huet F, Fry AM, Saunier S, Rivière JB, Attié-Bitach T, Garcia-Verdugo JM, Faivre L, Mégarbané A, Nachury M v. 2014. The oral-facial-digital syndrome gene C2CD3 encodes a positive regulator of centriole elongation. Nature Genetics 46:905–911. doi:10.1038/ng.3031

Tsai JJ, Hsu W bin, Liu JH, Chang CW, Tang TK. 2019. CEP120 interacts with C2CD3 and Talpid3 and is required for centriole appendage assembly and ciliogenesis. Scientific Reports 9. doi:10.1038/s41598-019-42577-0

Vaughan S, Gull K. 2016. Basal body structure and cell cycle-dependent biogenesis in Trypanosoma brucei. Cilia. doi:10.1186/s13630-016-0023-7

Vorobjev IA, Chentsov YS. 1980. The ultrastructure of centriole in mammalian tissue culture cells. Cell biology international reports 4:1037–1044. doi:10.1016/0309-1651(80)90177-0

Wang Lei, Failler M, Fu W, Dynlacht BD. 2018. A distal centriolar protein network controls organelle maturation and asymmetry. Nature Communications 9. doi:10.1038/s41467-18-06286-y

Wang L, Failler M, Fu W, Dynlacht BD. 2018. A distal centriolar protein network controls organelle maturation and asymmetry. Nature communications 9:3938. doi:10.1038/s41467-018-06286-y

Yamamoto S, Kitagawa D. 2021. Emerging insights into symmetry breaking in centriole duplication: updated view on centriole duplication theory. Current Opinion in Structural Biology. doi:10.1016/j.sbi.2020.08.005

Yang TT, Chong WM, Wang WJ, Mazo G, Tanos B, Chen Z, Tran TMN, Chen Y de, Weng RR, Huang CE, Jane WN, Tsou MFB, Liao JC. 2018. Super-resolution architecture of mammalian centriole distal appendages reveals distinct blade and matrix functional components. Nature Communications 9:1–11. doi:10.1038/s41467-018-04469-1

Ye X, Zeng H, Ning G, Reiter JF, Liu A. 2014. C2cd3 is critical for centriolar distal appendage assembly and ciliary vesicle docking in mammals. Proceedings of the National Academy of Sciences of the United States of America 111:2164–2169. doi:10.1073/pnas.1318737111

Yubuki N, Leander BS. 2013. Evolution of microtubule organizing centers across the tree of eukaryotes. The Plant journal : for cell and molecular biology 75:230–244. doi:10.1111/tpj.12145

Zhang D, Aravind L. 2012. Novel transglutaminase-like peptidase and C2 domains elucidate the structure, biogenesis and evolution of the ciliary compartment. Cell Cycle 11:3861– 3875. doi:10.4161/cc.22068

