## Supplemental Figure Legends for "Evolutionary conservation of centriole rotational asymmetry in the human centrosome"

**Supplemental Figure S1. Characterization of LRRCC1 expression in CRISPR or RNAi-treated cells. a)** LRRCC1 localization in non-treated RPE1 cells (left), or in cells treated with nocodazole to disperse pericentriolar satellites (right). Anti-LRRCC1 (Ab1, yellow), □-tubulin (magenta) and DNA (cyan). Bar, 5 µm (insets, 2 µm). **b)** Western blot analysis of over-expressed GFP-LRRCC1 fusions using anti-LRRCC1 (Ab2) or anti-GFP antibodies. Cell lysates were obtained from HEK 293 cells induced (+ Dox) or not (-Dox) to express GFP-LRRCC1 fusions in which the GFP is inserted either after aa 251 or 402. The load represents the same number of cells for all conditions. The different samples were deposited in duplicate and the labeling with the two antibodies was performed in parallel and exposed in the same way. Note that GFP fusions are already detected in the non-induced samples, whereas the endogenous protein (expected size ~120 kDa) is not. **c)** qRT-PCR analysis of *LRRCC1* expression in the CRISPR clones. mRNA levels are expressed as percentages of RPE1 control values. Bars, mean ± SD, 3 independent experiments. p values are provided when statistically significant from the corresponding control (One-way ANOVA). **d)** LRRCC1 centrosomal levels in CRISPR mutant cells stained with Ab1. Bars, mean ± SD, 3 independent experiments. p values are provided when statistically significant from the corresponding control (One-way ANOVA). **e)** LRRCC1 centrosomal levels in nocodazole-treated control or CRISPR mutant RPE1 cells stained with Ab2. Bars, mean ± SD, 3 independent experiments. p values are provided when statistically significant from the corresponding control (One-way ANOVA). **f)** Percentage of ciliated cells in serum deprived RPE1 cells treated with control or LRRCC1 siRNAs. Bars, mean ± SD, ≥ 83 cells from 3 independent experiments for each condition. p values are provided when statistically significant from the corresponding control (One-way ANOVA). **g)** LRRCC1 centrosomal levels in control or CRISPR edited (clone 25) HEK 293 cells stained with Ab1. Bars, mean ± SD, 3 independent experiments. p values are provided when statistically significant from the corresponding control (One-way ANOVA). **h)** Centriole and procentriole in a RPE1 cell processed for U-ExM and stained with anti-LRRCC1 (Ab1, yellow) and acetylated tubulin (magenta). Bar, 0.2 µm. **i)** qRT-PCR analysis of *PTCH1* expression in serum-deprived WT or CRISPR RPE1 cells treated with SAG or DMSO during 24 hours, expressed as percentages of the DMSO-treated control. Mean ± standard deviation, 2 independent experiments. **j)** Number of centrioles per spindle pole in WT mitotic RPE1 cells, CRISPR clones 1.1 and 1.9, or WT cells treated with control or LRRCC1 siRNAs during 48 hours. Centrioles were labelled with an antibody against hPOC5. Bars, mean ± SD, 50 cells from 3 independent experiments per condition. **k)** Duration of mitosis in WT RPE1 cells and in LRRCC1-deficient CRISPR clones. Bars, mean ± SD, 100 cells from 3 independent experiments. p values are provided when statistically significant from the corresponding control (One-way ANOVA).

**Supplemental Figure S2. Pipeline for generating average protein maps. a)** Examples of verticalized and annotated centrioles (one centriole per row). (A, B): XY cross section taken at z-position, at which the XY reference points have been provided. (C, D): YZ cross section taken at x-position, at which the centriole center has been calculated (from XY reference points). The mentioned z-position and x-position are shown with blue lines; red lines in the right columns show the Z reference points: the top and the bottom of the provided rectangular frame. Note that the centrioles significantly differ in their diameters and lengths (in pixels), and that the protein of interest is not always positioned in the same manner. **b)** Examples of standardized images. (A, B): XY cross section taken at a fixed z-position slightly under the top of the Z reference rectangle (note that this position is slightly different from that at which the XY reference points were provided). (C, D): YZ cross section taken in the middle of the XY plane (the standardized centrioles are always placed with their centers in the middle of the image). The mentioned z-position and x-position are shown with blue lines; red shapes outline a cylinder with the target standard sizes: diameter 0.8 µm (4 x expanded 0.2 µm), length 3 µm; image calibration 0.01 µm/pixel (all three parameters are tunable). Note that in contrast to the centrioles in a), the diameter and the length of the standardized centrioles and the position of the protein of interest in the XY plane are very similar.

**Supplemental Figure S3. Genomic characterization of the 1.1 CRISPR cell line and analysis of the corresponding transcripts. (a)** Scheme showing the deletions and insertions observed in the genome of the CRISPR 1.1 line. In one copy of the *LRRCC1* locus (Chromosome 8.a), a 179 bp fragment corresponding to part of the deleted sequence is inserted in antisense orientation. Exons are represented by dark grey boxes and are numbered. **(b)** Comparison of wild type and 1.1 transcripts. Top: Wild type LRRCC1 isoforms. Only the splicing of exon 2 is validated by comparison with EST databases. The location of the Leucine Rich Repeat (LRR) and coiled-coil domains is indicated. Bottom: Two transcripts were detected in the CRISPR line 1.1. Isoform 1a exhibits splicing of exons 7-8 and is thus transcribed from chromosomes 8.a, and isoform 1b exhibits splicing of exons 9-10 and thus transcribed from chromosomes 8.b. Both transcripts are in frame but carry deletions compared to wild-type isoforms. Note that a fraction of these transcripts could also have a deletion of exon 2, as in wild type cells.

**Supplemental Figure S4. Quantification of DA or distal centriole components in LRRCC1-deficient cells.** Centrosomal levels of CEP164 (a, d), CEP290 (b, e), and OFD1 (c, f) in RPE1 CRISPR clones (a-c) and RNAi-treated RPE1 cells (d-f). Bars, mean ± SD, 3 independent experiments. p values are provided when statistically significant from the corresponding control (One-way ANOVA).

**Supplemental Figure S5. LRRCC1 does not interact directly with C2CD3.** Co-immunoprecipitation experiments from a lysate of HEK 293 cells expressing LRRCC1 with GFP inserted after aa 402. Anti-GFP or control (anti-HA tag) antibodies were used for immunoprecipitation, and Western blot was performed using either anti-GFP or anti-C2CD3 (RRID:AB_2718714) antibodies. Lys = lysate; SN = supernatant; IP = immunoprecipitation. The amount of lysate loaded on the gel represents 4% of the amount used for the immunoprecipitation.
