## Supplementary figures and images for "Evolutionary conservation of centriole rotational asymmetry in the human centrosome"

### Supplemental Figure S1

#### Figure S1

Gaudin *et al.*

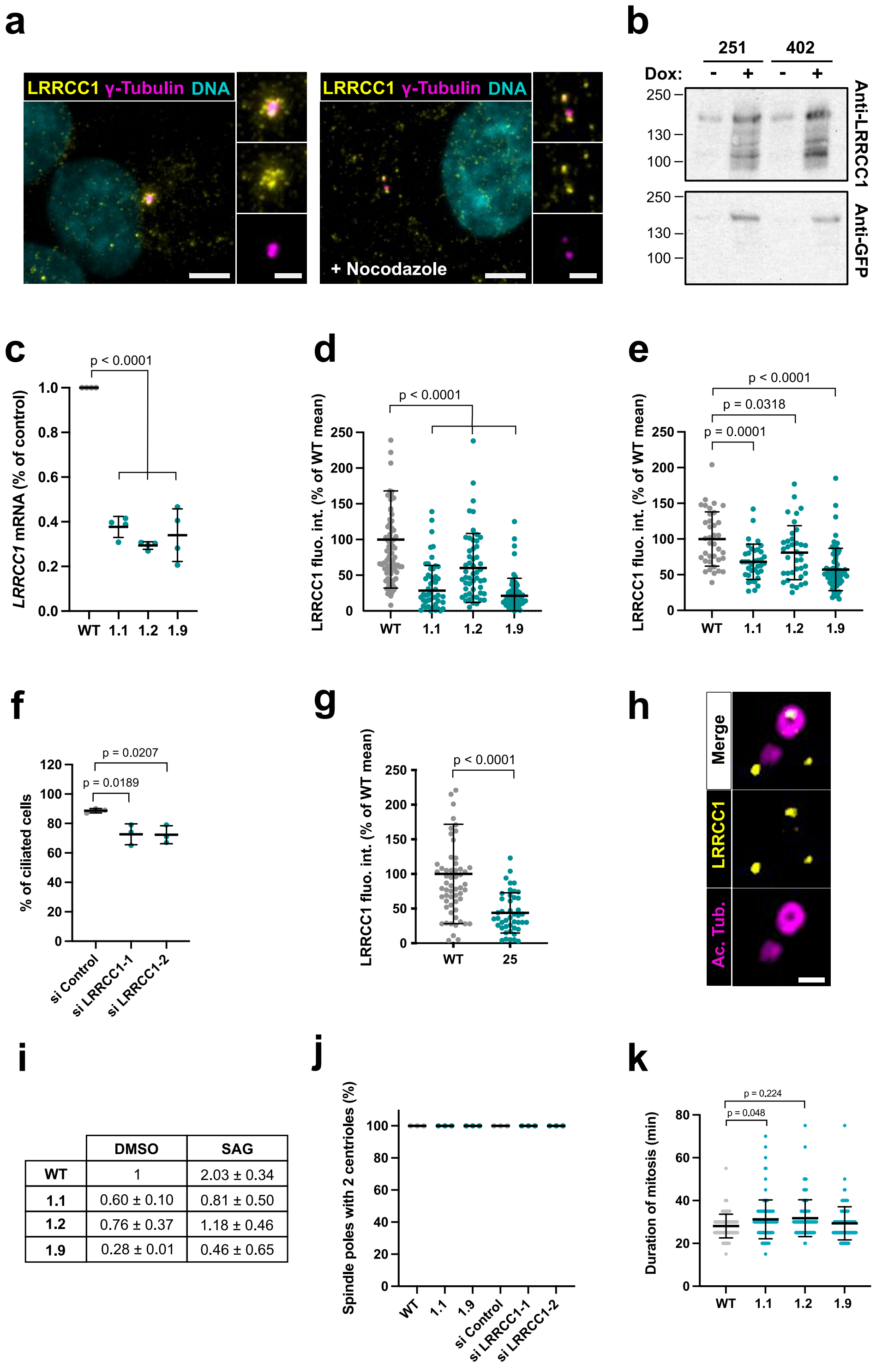


| | DMSO | SAG |
|---|---|---|
| WT | 1 | 2.03 ± 0.34 |
| 1.1 | 0.60 ± 0.10 | 0.81 ± 0.50 |
| 1.2 | 0.76 ± 0.37 | 1.18 ± 0.46 |
| 1.9 | 0.28 ± 0.01 | 0.46 ± 0.65 |

### Supplemental Figure S2

#### Figure S2

Gaudin *et al.*

**a**

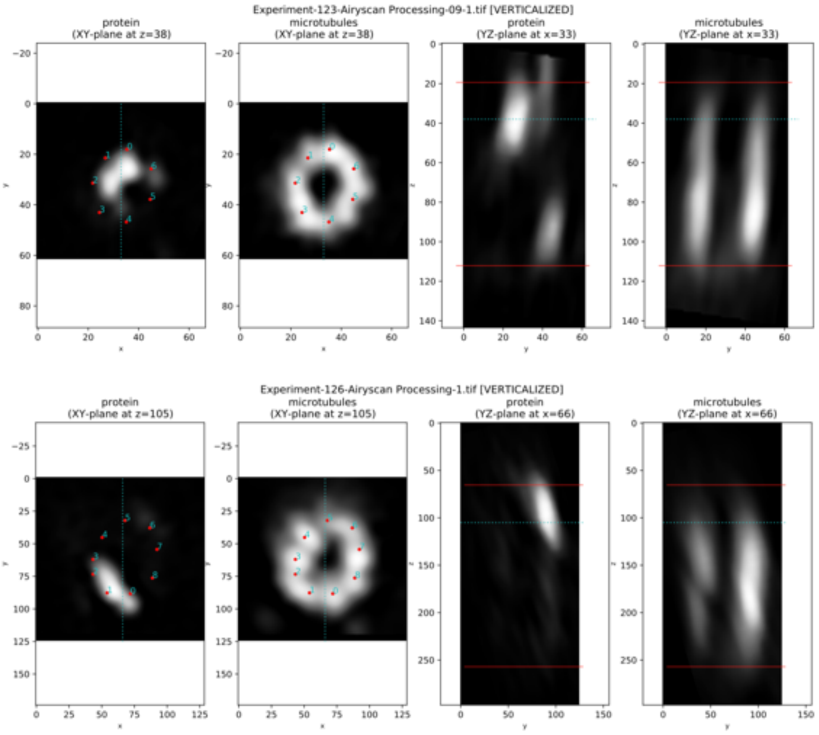


**b**

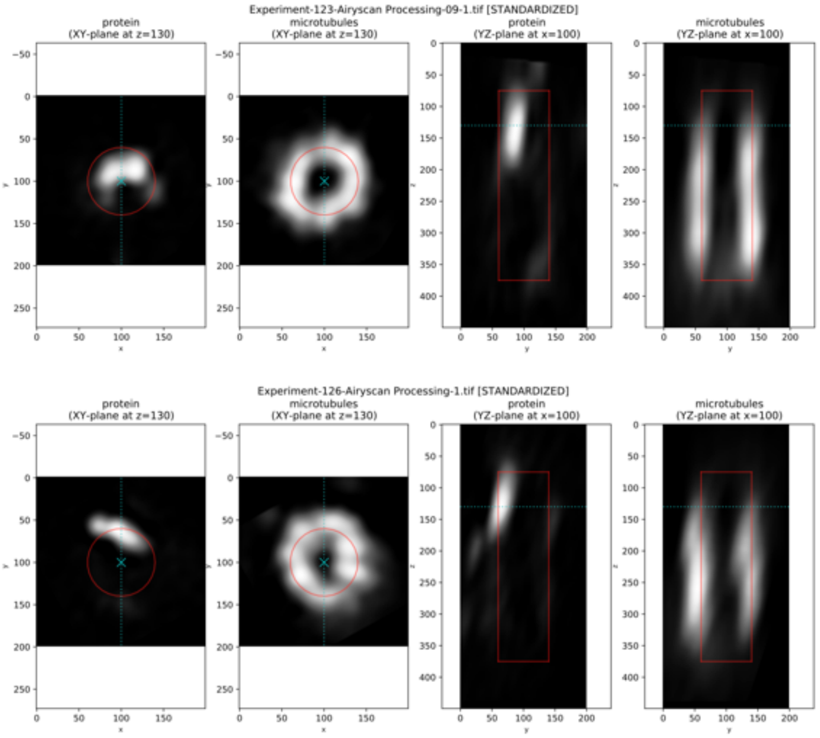

### Supplemental Figure S3

#### Figure S3

Gaudin *et al.*

## a

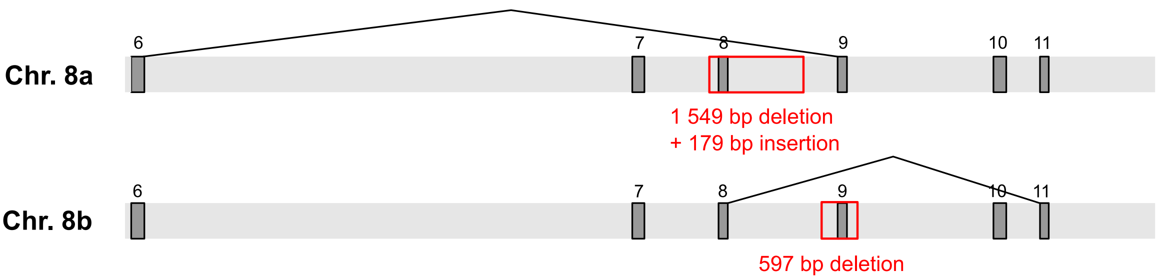


## b

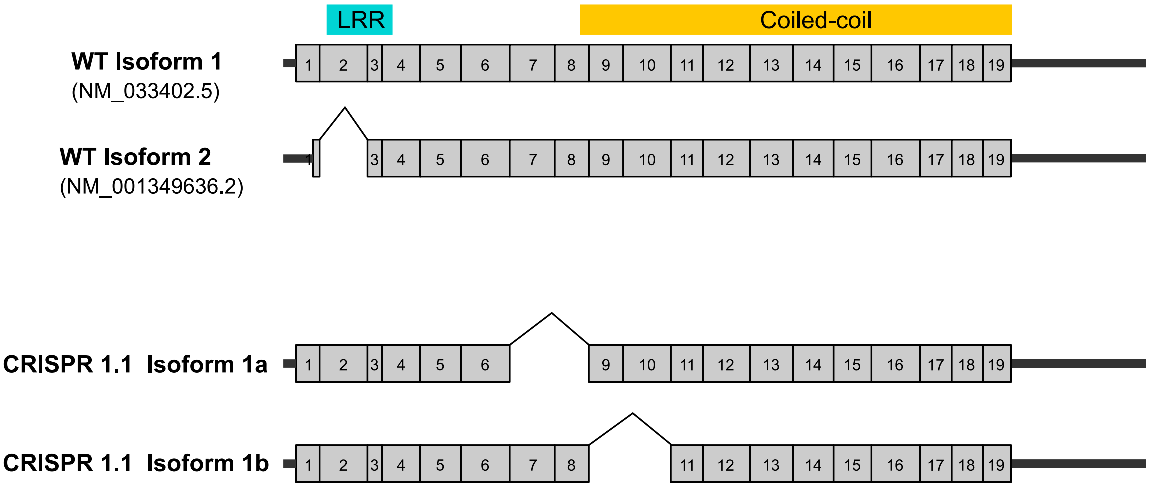

### Supplemental Figure S4

#### Figure S4

Gaudin *et al.*

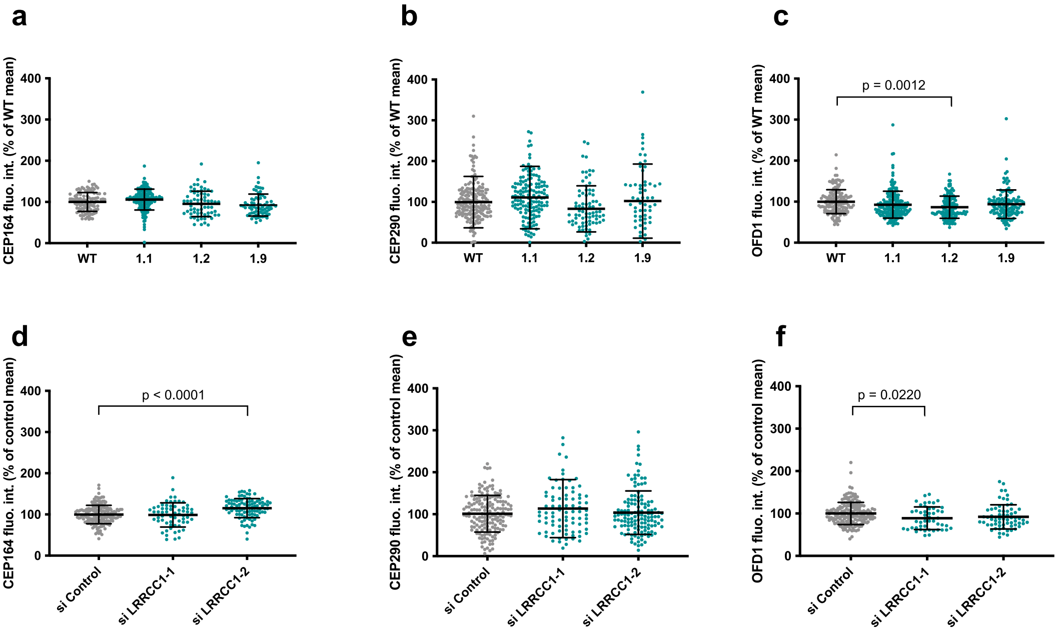

### Supplemental Figure S5

#### Figure S5

Gaudin *et al.*

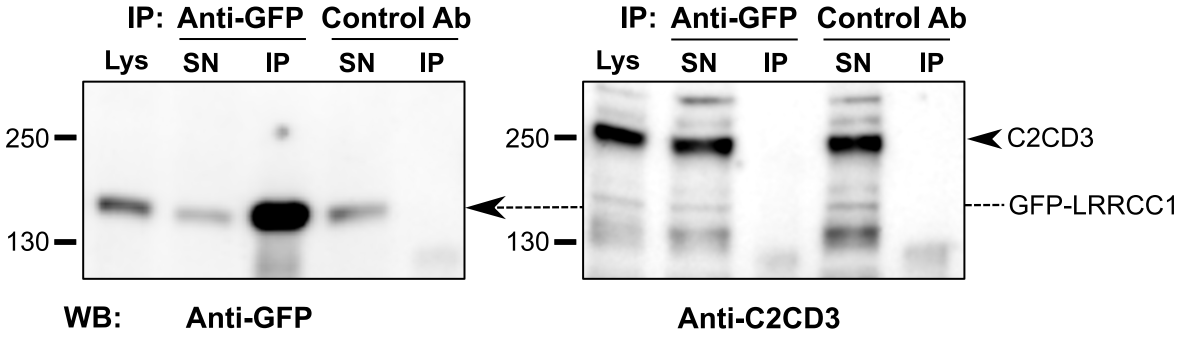
